# The effect of aging on post-translational modifications of wild-type human SOD1 and the A5V ALS mutant

**DOI:** 10.1101/2025.06.02.657411

**Authors:** Gabriel Freitas de Souza, Rayne Stfhany Silva Magalhães, José Raphael Monteiro Neto, Michele Martins, Cristian Follmer, Magno Junqueira, Elis Cristina Araujo Eleutherio

## Abstract

Cu/Zn superoxide dismutase 1 (SOD1) is essential for maintaining neural health. Its functions include modulating metabolism, maintaining redox balance, regulating transcription, besides eliminating superoxide radicals, which are achieved through various post-translational modifications (PTMs). Consequently, unusual PTMs in SOD1 can impair its functionality and stability, leading to the accumulation of misfolded SOD1 and the increase of oxidative stress markers, hallmarks of Amyotrophic Lateral Sclerosis (ALS). Although SOD1 has been extensively studied, especially regarding its role in ALS, relatively little is known about how aging and mutations affect SOD1 PTMs. This study aimed to evaluate the effect of oxidative stress induced by chronological aging on PTMs of human SOD1: wild-type (WT) and A5V SOD1, a severe ALS-related mutant. To do this, both hSOD1 forms were expressed in *Saccharomyces cerevisiae* lacking the SOD1 gene, and then purified from extracts of stressed and non-stressed cells. PTMs were analyzed using mass spectrometry, observing the modification of WT and mutant human SOD1 in both conditions. We observed changes in the levels of damage, including oxidation, formylation, and carboxylation, such as oxidized tryptophan 33, associated with prion-like propagation of SOD1 misfolding. Increased levels of this PTM appeared in WT SOD1 after aging and in A5V SOD1. Acetylation and succinylation were also found on lysines. Some of these modifications already have described functions in the literature, while others still lack a defined role. Interestingly, the levels of these physiological PTMs differed between WT and mutant SOD1, providing important information for elucidating the molecular mechanisms of ALS involving SOD1.

## Introduction

As people live longer, more research is exploring how aging at the cellular level contributes to diseases associated with aging, such as neurodegenerative, metabolic, cardiovascular, and bone diseases. Recent studies have pinpointed indicators of cellular aging; for example, Lopez-Otín and colleagues outlined the key features of aging and health [1]. They suggested that oxidative stress is involved in several biomarkers of cellular aging, including mitochondrial dysfunction, loss of proteostasis, accumulation of misfolded proteins, and alterations in epigenetic control [2–3].

Post-translational modifications (PTMs) may connect oxidative stress to markers of cellular aging [2]. PTMs of proteins often increase their stability, control their activity, and affect where they are located within the cell. Furthermore, because PTMs affect different tissues and organs in specific ways, research indicates they play a significant role in the development of diseases. These include the age-related diseases such as Parkinson’s disease, Huntington’s disease, and amyotrophic lateral sclerosis (ALS) [2].

ALS is a rare, rapidly progressing, and well-studied neurodegenerative disease. It affects both upper and lower motor neurons, which disrupts muscle control. At a molecular level, ALS involves the loss of specific nerve cells that show a buildup of misfolded proteins and signs of oxidative stress [4]. About 90% of ALS cases occur randomly (sALS), while the rest are inherited (fALS) [5]. Regardless of the type, ALS advances quickly after symptoms begin, and the typical life expectancy is 2 to 5 years. PTMs may contribute to the factors driving this rapid progression and could potentially connect sALS and fALS. Research indicates that PTMs are found in samples from both forms of the disease [6].

One of the most studied biomarkers in ALS is the SOD1 proteinopathy [7]. SOD1, a 32 kDa homodimer composed of 153 amino acids, is crucial for neural healthy. It eliminates superoxide radicals in the cytoplasm and intermembrane mitochondrial space (IMS), activates antioxidant pathways when in the nucleus, and regulates the mitochondrial electron transport chain’s respiratory rate [5]. Over 200 mutations in the SOD1 gene have been linked to inherited ALS [8]. Most mutations cause a specific amino acid substitution, while others result in deletions, insertions, truncations, or changes in the reading frame during translation [9, 10].

The functions of SOD1 depend on its location within the cell and are modified by different PTMs [5,6]. For example, acetylation at Lys71 causes SOD1 to detach from its copper chaperone (CCS). This separation allows SOD1 to relocate to IMS, where it helps maintain superoxide balance [11]. Within this mitochondrial compartment, SOD1 not only eliminates superoxide but also regulates its production by modulating Complex I activity through succinylation at Lys123 [12]. Phosphorylation at Ser60 and 99, as well as palmitoylation at Cys7, induce SOD1 translocation to the nucleus, where it activates oxidative stress response [13, 14]. In the cytosol, SOD1 activity is modulated by nutrient availability via phosphorylation at Thr40 mediated by mTORC1 [15]. SOD1 also participates in the regulation of aerobic glycolysis by influencing pyruvate dehydrogenase phosphorylation [16]. This regulation is particularly important in astrocytes because it allows them to provide lactate to neurons. This, in turn, enables neurons to conserve glucose, which they can then use for NADPH production to protect against oxidative stress.

For SOD1 to function properly, it must fold into its correct three-dimensional structure. This folding process relies on key modifications. First, a disulfide bond forms within the SOD1 molecule, specifically between Cys58 and Cys147. This bond stabilizes the individual SOD1 unit and helps it fold correctly. Second, copper and zinc ions must bind to SOD1, which is essential for its enzymatic activity and structural stability. Third, a hydrophobic region of SOD1 needs to be exposed, allowing it to pair up with another SOD1 unit to form a dimer [17]. When these modifications are disrupted — for example, by SOD1 mutations or damaging modifications caused by oxidative stress — SOD1 can misfold and lose its function [17]. Moreover, oxidative changes to SOD1 related to aging are thought to not only cause SOD1 to clump together, but also to promote the creation and buildup of other harmful SOD1 forms, such as monomers, trimers, and oligomers. These factors may speed up cellular aging in ALS, as well as in other neurodegenerative diseases such as Alzheimer’s and Parkinson’s [13].

The role of PTMs in the development and progression of disease has been well-documented. Bedja-Iacona and colleagues examined tissues from deceased ALS patients and identified differing amounts of 15 types of PTMs at 14 locations on the SOD1 protein [18]. These differences were found in 69% of the ALS patients they studied. Additionally, it is believed that SOD1 clumps in sALS cases may be related to PTMs resulting from oxidative stress, including the oxidation of specific cysteine and histidine amino acids [14]. Therefore, identifying PTMs in target proteins associated with aging is important for understanding the causes of age-related diseases. Techniques like flow cytometry, protein microarrays, Western blotting, and enzyme-linked immunosorbent assays (ELISA) can be used to study PTMs [19]. While these methods are well-established, they have limitations, such as the limited specificity and selectivity of the antibodies used [20]. In contrast, mass spectrometry has become a primary technique for studying PTMs because of its speed, resolution, and ability to identify and measure PTMs in complex protein mixtures [20]. Furthermore, by combining databases and bioinformatics analyses with mass spectrometry, we can precisely locate modifications on proteins, which helps us understand how these modifications affect protein function and stability [21].

Recent research has increasingly focused on understanding the mechanisms of PTMs and their role in the progression of ALS. These studies aim to develop therapies that prevent harmful PTMs, especially those linked to oxidative stress. Therefore, this study aimed to investigate and quantify the PTM profiles of wild-type (WT) SOD1 and A5V SOD1 related to cellular aging. This SOD1 mutation is one of the most aggressive. Individuals with the A5V mutation can develop the disease at any age, with an average survival of only 1.4 years after the first symptoms [22]. Phenotypically, the A5V mutation is marked by a sudden and marked onset of symptoms, with a very rapid progression of the disease, usually manifesting as muscle weakness in the limbs or bulbar muscles [10].

## Results

### Purification procedure enriches SOD1 in all samples

To determine the extent of enrichment of WT and mutant SOD1 obtained from control and aged cells, we measured SOD1 enzymatic activity before and after purification process (Figures S1 and 2a). Purification led to roughly a 2- to 3-fold increase in activity for both WT and mutant SOD1 under both conditions. This suggests that the purified samples had a greater proportion of SOD1 compared to the crude extracts, indicating effective enrichment of the protein. This result also suggests that the SOD1 purification method did not negatively affect the protein’s structure PTMs required for its enzymatic activity (binding to Cu^2+^ and Zn^2+^ ions). The previously observed decrease in A5V SOD1 activity compared to WT has been reported by our group [16].

### Aging and A5V mutation affect the carbonylation and phosphorylation profile of SOD1

To evaluate the levels of SOD1 carbonylation and phosphorylation in the enriched extracts, specific antibodies targeting DNP (carbonyl) or phosphorylated serine (pSer) were used (Figure 2b-c). Cabonylation levels of WT SOD1 increased after aging, while in the mutant SOD1, they did not change (Figure 2b). Carbonylation is associated with oxidative stress and may lead to protein dysfunction due to the oxidation of the side chain of lysines, histidines, and cysteines to ketones or aldehydes.

**Figure 1.**
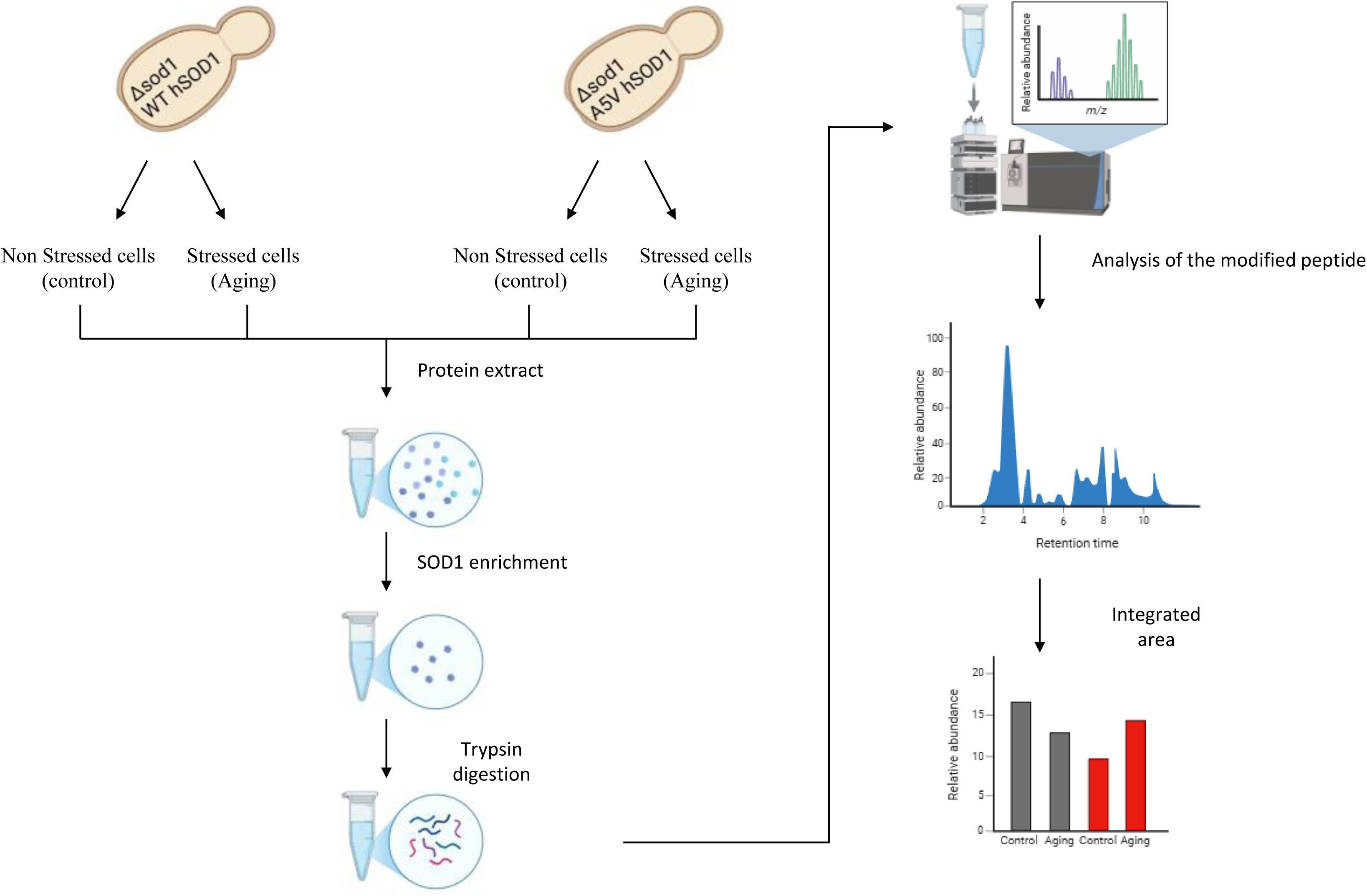
Workflow for the identification and quantification of PTMs on human SOD1 (hSOD1) in response to oxidative stress. hSOD1 was obtained from cellular extracts of humanized yeast cells, where their endogenous SOD1 was replaced by WT hSOD1 or A5V mutant. Yeast cells grown in glucose obtain energy solely through fermentation and are therefore not exposed to oxidative stress, serving as a control condition. These cells were then transferred to water (a non-proliferative condition) and incubated at 37 °C/160 rpm during 24 h. The combination of high temperature and aeration induces substantial oxidative stress, speeding up the aging process. Protein extracts were enriched with hSOD1 and digested with trypsin. Peptides were eluted and analyzed by LC–MS-MS. Proteomics data were analyzed with Proteome Discoverer 2.1 software and Xcalibur 4.6. Peptides were quantified by precursor ion area detection and normalized through peak integration of a proteotypic peptide.

**Figure 2.**
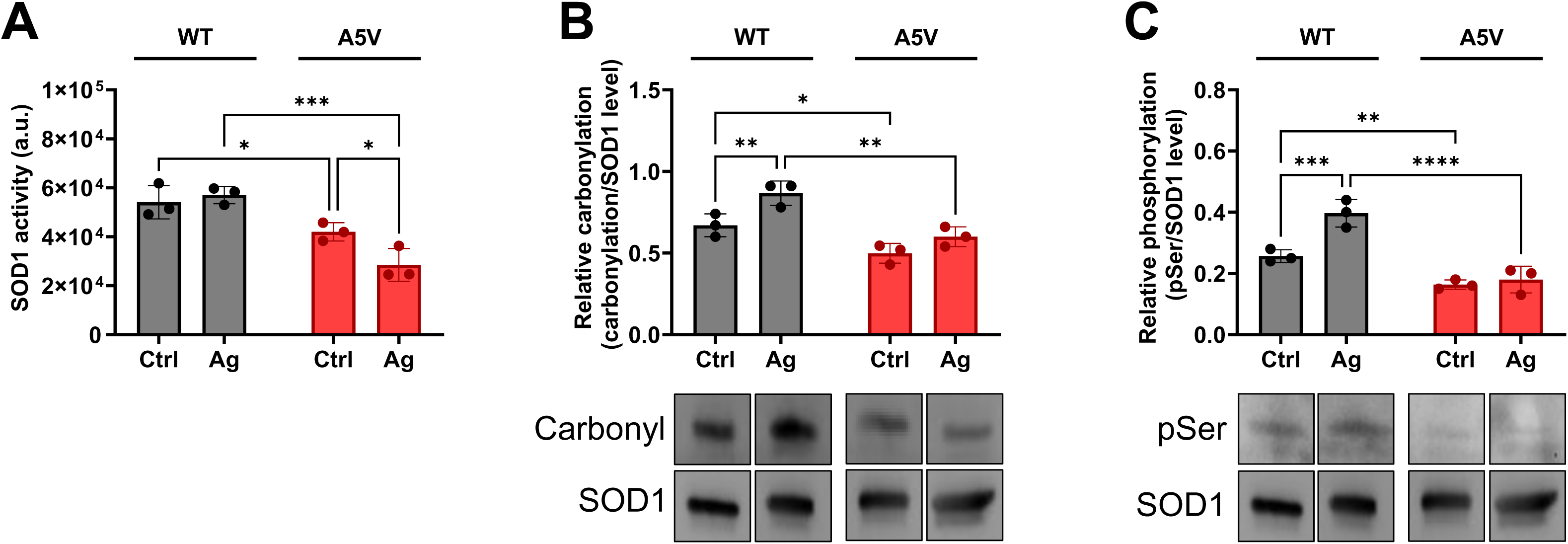
Effect of aging on enzymatic activity, carbonyl, and pSer levels of WT and A5V SOD1. Cellular extracts were obtained from cells grown under fermentative conditions, which resulted in low levels of reactive oxygen species - ROS (Ctrl), and after chronological aging (Ag), which increases oxidative stress. These extracts were subjected to protein purification methods to enrich the samples with human SOD1. (A) SOD1 activity was quantified after enrichment; (B) Levels of carbonyl and (C) pSer were determined using immunoblotting assays and normalized to total SOD1 levels. Data are expressed as Mean ± SD from at least three independent experiments. Two-way ANOVA was used for statistical analysis with a significance level of * p < 0.05, ** p < 0.01, *** p < 0.001, **** p < 0.0001.

Another PTM analyzed was pSer. Previous large-scale data analyses indicate that human SOD1 can be phosphorylated at eight serine residues: Ser26, Ser35, Ser60, Ser69, Ser99, Ser103, Ser106, and Ser108 (https://www.phosphosite.org/). The roles of most of these phosphorylation sites remain unclear, except for positions 60 and 99, which have been linked to SOD1 transport to the nucleus, where SOD1 activates the expression of genes involved in protection against oxidative stress [26]. Our results showed an approximate 1.5-fold increase in pSer levels in WT SOD1 after aging, while levels in the mutant remained unchanged (Figure 2c). This suggests that A5V SOD1 is not sensitive to oxidative stress as a regulator of nuclear translocation, which supports earlier findings from our group [27]. However, it is important to notice that this result could be related to the phosphorylation of any serine residue in SOD1.

### Scanning of the SOD1 sequence and identification of a normalizing peptide

Next, we used MS to identify and quantify modified residues of WT and mutant SOD1 obtained from cells submitted to low or high ROS concentrations. Immunoblotting and MS are both used to identify PTMs. A limitation of immunoblotting is the need for specific antibodies to detect particular PTMs, which MS does not require. Moreover, MS is generally more accurate and sensitive for identifying PTMs, especially when characterizing novel or site-specific modifications.

To determine sequence coverage, total MS/MS spectra, the number of unique peptides, and to extract all XICs of both unmodified and modified peptides of SOD1, the raw were manually inspected and analyzed using Xcalibur 4.6 and Proteome Discoverer 2.1, respectively. A greater number of unique peptides analyzed from a protein increases the probability of obtaining more accurate results regarding modifications. Trypsin digestion produced nineteen SOD1 peptides, all of which were detected during the scanning process (Table S1). Furthermore, all the peptides were considered unique to SOD1. This equivalence between the total number of peptides found and the total number of unique SOD1 peptides is an important factor. It ensures that the peptide selected for standardizing chromatogram areas is a proteotypic SOD1 peptide and that all peptides analyzed by manual integration of peak areas in the chromatogram are solely related to SOD1, reducing the risk of false positives. The analysis also revealed 93.50649% coverage of the total SOD1 sequence, indicating minimal peptide loss during digestion. This suggests that nearly the entire SOD1 sequence was analyzed, minimizing the possibility of missing any PTMs.

Of the nineteen unique SOD1 peptides identified in both WT and A5V SOD1, twelve showed PTMs. Analysis of the chromatograms revealed six different modifications: acetylation, phosphorylation, succinylation, oxidation, formylation, and carboxylation (Table 1). These modifications can be classified into two groups: (a) physiological modifications, which regulate SOD1’s function, structure, and localization. Acetylation, succinylation, and phosphorylation belong to this group; and (b) modifications resulting from cellular stress, such as oxidative stress caused by cellular aging. These stress-related modifications can damage SOD1, leading to dysfunction, structural disruption, and mislocalization of the protein. Formylation, oxidation, and carboxylation fall into this second group.

**Table 1.** Modifications found in human SOD1 from the analysis of data obtained through MS and data processing in the Proteome Discoverer 2.1 software. The modifications found in each SOD1 residue, physiological or damage, were listed.

| **PTM in SOD1** | **Amino Acid** | **Position in the protein** |
| --- | --- | --- |
| **Formylation** | Thr | 55, 59, 89, 136 |
|  | Arg | 70, 116 |
|  | Lys | 92, 137 |
| **Carboxylation** | Thr | 59, 89 |
|  | Arg | 116 |
|  | Lys | 92, 129, 137 |
| **Acetylation** | Lys | 10, 92, 123, 129, 137 |
| **Oxidation** | Trp | 33 |
|  | Lys | 4, 37 |
|  | His | 47, 49, 81, 111, 121 |
|  | Asn | 87, 132, 140 |
| **Succinylation** | Lys | 10, 137 |
| **Phosphorylation** | Thr | 3, 55 |

Manual integration was performed using Xcalibur 4.6 to quantify PTMs in each peptide. This involved integrating the area under the curve in the chromatogram of the modified peptide, filtering by the m/z ratio, and identifying the retention time of each modified peptide. Furthermore, normalization was performed using a non-modified SOD1 proteotypic peptide present in all samples, based on the raw data. For this normalization, four different peptides were considered as potential normalizers. Initially, the relationship between the intensity of the peptides selected for normalization and the peptide spectrum match (PSM) parameter, which represents the result of matching a peptide to an experimental mass spectrum, was examined. We confirmed that all the chosen peptides showed a tendency for their intensity to increase with increasing PSM values, demonstrating a good correlation between peptide area and PSM values in the samples (Figure S2). A Pearson correlation analysis was performed to visualize the consistency of normalized modifications within each condition and its triplicate (Figure S3). We confirmed that the normalized data exhibited a good correlation, indicating that the selected peptides did not negatively impact the correlation between the biological samples. In addition, a principal component analysis (PCA) was performed to observe how normalization affects the clustering of the biological triplicates (Figure S4). The peptide [K].HGGPKDEER.[H] (m/z 512.74456) resulted in a better clustering of the biological replicates, suggesting that it was the most suitable peptide for the normalization of raw data.

### Physiological PTMs were altered by aging and the A5V mutation

According to PhosphoSitePlus (https://www.phosphosite.org/), lysine residues of human wild-type (WT) SOD1 can be both acetylated and succinylated at four positions (10, 92, 123, and 137), solely acetylated at three positions (4, 24, and 71), and solely succinylated at two positions (76 and 129). Our results showed that both WT and A5V mutant SOD1 were acetylated at lysine residues 10, 92, 123, 129, and 137 (Figure 3a-e). Lys137 was also found to be succinylated (Figure 3f). The literature indicates that lysine succinylation is a common modification that often overlaps with acetylation [28]. In both WT SOD1 and A5V SOD1, oxidative stress appeared to decrease the level of SOD1 peptides acetylated at lysines 10, 92, and 123. However, the levels of these modified peptides were higher in mutant SOD1 than in WT under control conditions.

**Figure 3.**
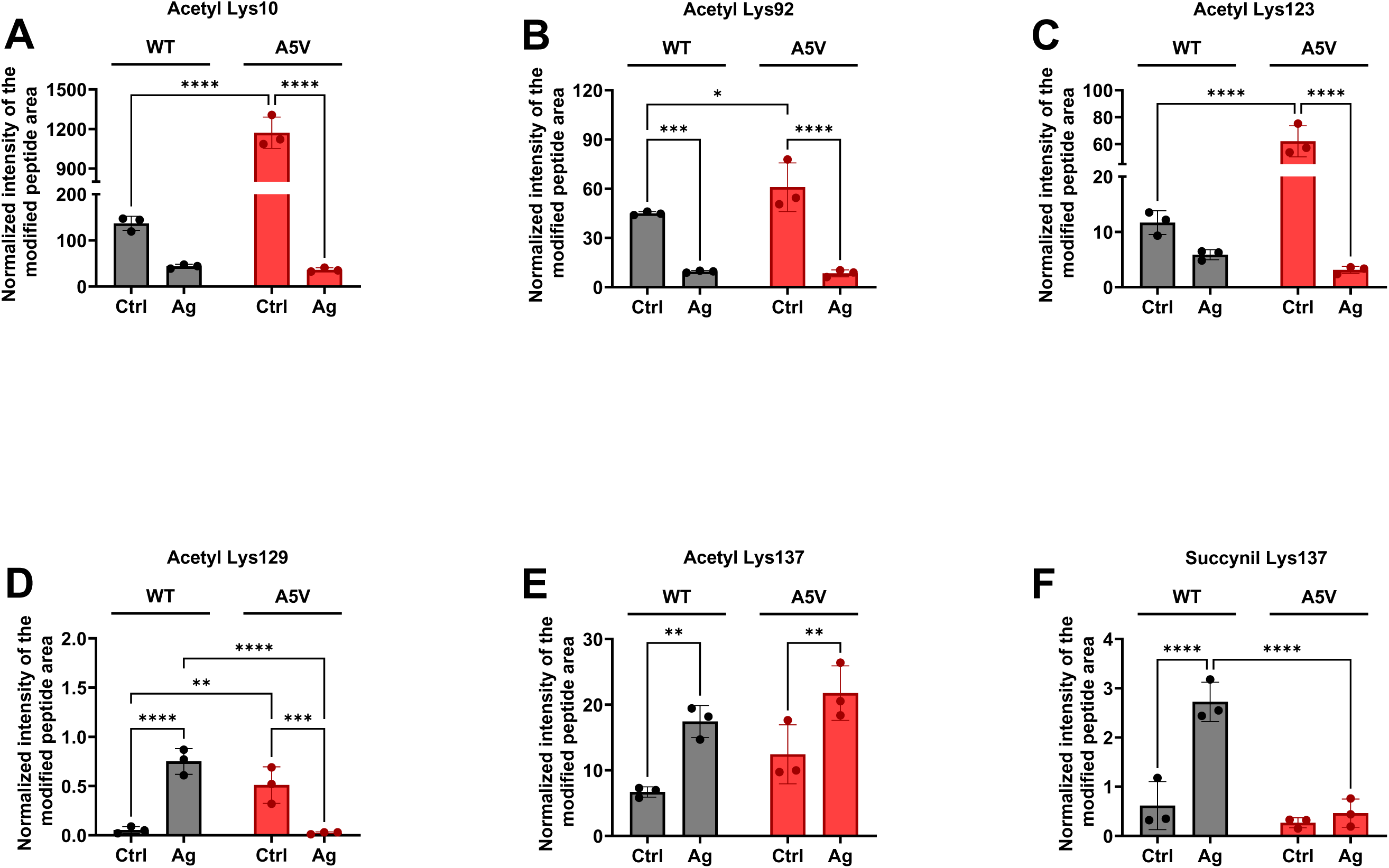
Aging modifies the profile of physiological modifications in lysines of WT and A5V SOD1. Cellular extracts were obtained from non-stressed cells (Ctrl) and following chronological aging (Ag). Protein extracts enriched for SOD1 were processed using MS. The raw data were analyzed with Proteome Discoverer 2.1 and Xcalibur 4.6, and peptides were quantified by precursor ion area detection, normalized by peak integration of the proteotypic peptide [K].HGGPKDEER.[H]. PTM quantification was performed for (A) Acetyl Lys10, (B) Acetyl Lys92, (C) Acetyl Lys123, (D) Acetyl Lys129, (E) Acetyl Lys137, and (F) Succinyl Lys137. Data were expressed as Mean ± SD from at least three independent experiments. Two-way ANOVA was applied for statistical analysis with significance levels of * p < 0.05, ** p < 0.01, *** p < 0.001, **** p < 0.0001.

The acetylation levels of Lys10 decreased after aging in the WT form, although this change was not statistically significant (p = 0.0921). In contrast, the mutant form showed a significant reduction in acetylation of approximately 100-fold after chronological aging (Figure 3a). Under control conditions, acetylation levels of Lys10 in the mutant strain were higher than those observed in both conditions of the WT strain (Figure 3a). The acetylation levels of Lys92 in WT decreased approximately four-fold in aged cells (Figure 3b). Currently, no function has been described in the literature for the acetylation of SOD1 at lysines 10 and 92. This modification could be a strategy to reduce SOD1 aggregation propensity, which is increased in the A5V mutant [27]. Acetylation changes the net charge of lysine from +1 to 0, increasing the net negative charge of SOD1, which may protect against proteinopathy [28].

Statistical analysis revealed that the levels of acetylated Lys123 within the SOD1 peptide remained constant in WT SOD1 following oxidative stress. However, these levels decreased approximately six-fold in the mutant SOD1. Literature suggests that deacetylation at Lys123 allows SOD1 to suppress respiration, contributing to SOD1-mediated antioxidant defense [29]. The analyses also unexpectedly showed acetylation at Lys129 of SOD1. In WT SOD1, acetylation of this residue increased approximately seven-fold after cell aging, while the mutant SOD1 exhibited higher levels under control conditions compared to both aged and control WT SOD1.

Our data indicated that succinylation at Lys137 of WT SOD1 increased approximately three-fold after chronological aging. In contrast, the levels in the A5V mutant remained unchanged and lower than those in the WT form. We also detected SOD1 succinylation at Lys10; however, this modification was found in the peptide [1].MATKAVCVLKGDGPVQGIINFEQK.[26] along with the oxidation of Lys4. Therefore, we cannot definitively attribute the observed levels solely to succinylation. The role of succinylation at these SOD1 residues has not yet been explored in the literature. It may protect against protein aggregation, similar to what has been suggested for acetylation PTM. Succinylation may be more effective than acetylation in preventing SOD1 aggregation, given the greater negative charge gained by succinylated lysine (change from +1 to -1).

SOD1 peptides phosphorylated at Thr3 and Thr55 were also detected. However, these peptides contained other modifications in addition to phosphorylation, preventing quantification of phospho-Thr.

### Aging increases damaging modifications in the WT strain

The presence of oxidation, or hydroxylation as it is also known, is a marker of oxidative stress. The strategy addressed in our work for cellular aging leads to an increase in oxidative stress, as demonstrated in previous works by our group [24]. Here, we show that the oxidation levels of residues Trp33 (Figure 4a), His47 (Figure 4c), and His111 (Figure 4h) of WT SOD1 increased after aging. The opposite effect was noted in the His49 residues of WT SOD1 (Figure 4e). However, we have identified the presence of this modification in another peptide, accompanied by additional damaging modifications (Figure 4d). The oxidation levels of Lys37 (Figure 4b) and Asn87 (Figure 4g) residues were not significantly altered compared to the control condition but showed a tendency to increase. Moreover, an increase in oxidation of His81 and His111 was observed in association with the carboxylation of Thr89 and Arg116 within the same peptide (Figure 3f).

**Figure 4.**
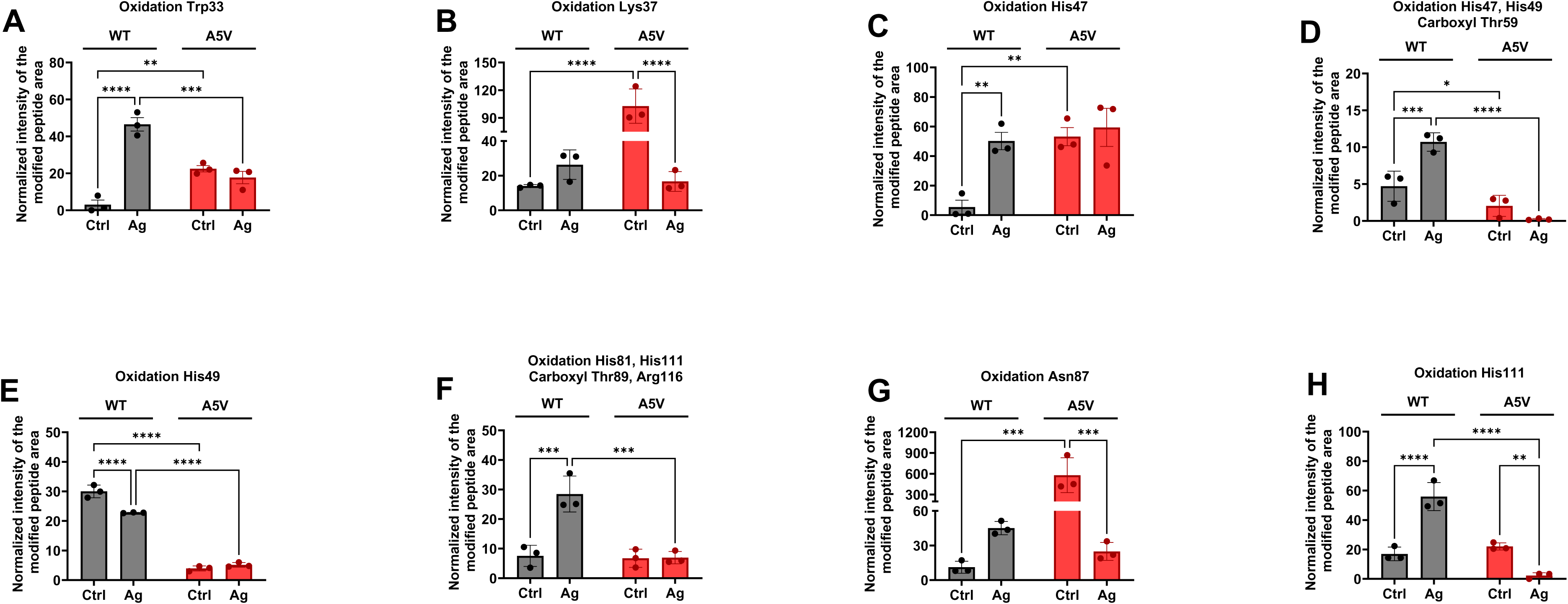
Oxidative damage in WT SOD1 residues exhibited greater sensitivity to aging compared to the A5V SOD1 mutant. Cellular extracts were obtained from non-stressed cells (Ctrl) and during chronological aging (Ag). Protein extracts enriched for SOD1 were processed using MS. The raw data were analyzed with Proteome Discoverer 2.1 and Xcalibur 4.6, and peptides were quantified by precursor ion area detection, normalized using peak integration of the proteotypic peptide [K].HGGPKDEER.[H]. Oxidative modifications were observed and quantified in residues (A) Trp33, (B) Lys37, (C) His47, (D) His47 and His49, alongside Thr59 carboxylation, (E) His49, (F) His81 and His111, alongside Thr89 and Arg116 carboxylation, (G) Asn87, and (H) His111. Data were expressed as Mean ± SD from at least three independent experiments. Two-way ANOVA was applied for statistical analysis with significance levels of * p < 0.05, ** p < 0.01, *** p < 0.001, **** p < 0.0001.

Carboxyl groups were also identified at residues Thr59, Lys92, Lys129, and Lys137. In WT SOD1, the levels of carboxyl groups at Lys92 and Lys137 increased after aging (Figure 5b and c), while the carboxylation of Thr59 decreased (Figure 5a). However, carboxylated Thr59 in WT SOD1 was observed alongside other oxidative damage in Figure 4d, which increased in aged cells. The observation of carboxylation at Lys129, along with additional physiological modifications within the same peptide, complicated its quantification, as we cannot solely attribute the increase to carboxylation or damage.

**Figure 5.**
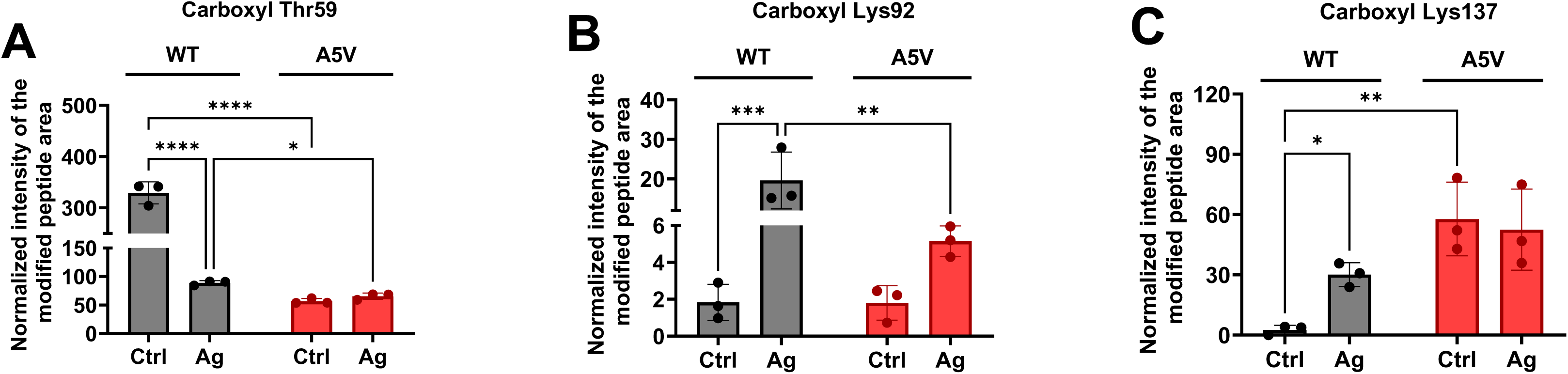
Carboxylation levels are differently affected by aging and the A5V mutation in SOD1. Cellular extracts were obtained from non-stressed cells (Ctrl) and during chronological aging (Ag). Protein extracts enriched for SOD1 were processed using MS. The raw data were analyzed with Proteome Discoverer 2.1 and Xcalibur 4.6, and peptides were quantified by precursor ion area detection, normalized through peak integration of the proteotypic peptide [K].HGGPKDEER.[H]. Carboxyl levels were quantified in residues (A) Thr59, (B) Lys92, and (C) Lys137. Data were expressed as Mean ± SD from at least three independent experiments. Two-way ANOVA was applied for statistical analysis with significance levels of * p < 0.05, ** p < 0.01, *** p < 0.001, **** p < 0.0001.

The addition of formyl groups increased in the WT SOD1 at residues Thr55 (Figure 5a), Arg70 (Figure 6c), Thr89 (Figure 6d), Arg116 (Figure 6f), and Lys137 (Figure 6h) after aging. The residues Thr59 (Figure 6b) and Lys92 (Figure 6e) did not show alterations in formylation levels under aging conditions. The formylation of residues Thr136 and Lys137 was observed to occur within the same peptide and exhibited an increase in the aged WT SOD1 compared to the control group (Figure 6g). Together, our observations indicate that most of the damaging modifications detected in WT SOD1 increased with chronological aging (Figure 7a). This aligns with previous findings, suggesting that SOD1 undergoes increased damage due to elevated oxidative stress associated with aging. These oxidative damages can affect the role of SOD1 (for example, oxidation of His 47 impairs Cu^2+^ link and, consequently, antioxidant activity) or promote SOD1 aggregation (for example, Trp33 oxidation).

**Figure 6.**
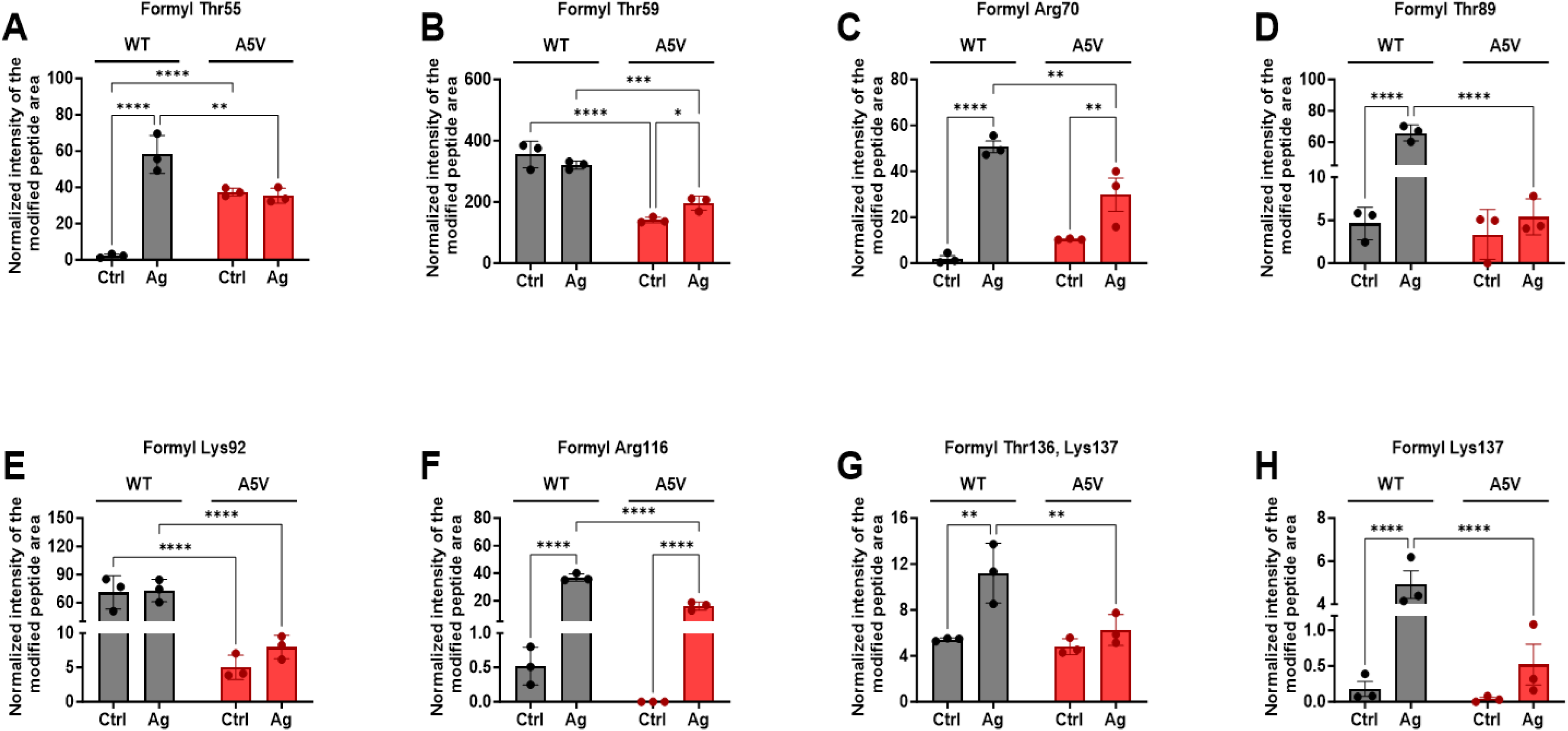
Formylation increases with aging in WT and A5V SOD1. Cellular extracts were obtained from non-stressed cells (Ctrl) and during chronological aging (Ag). Protein extracts enriched for SOD1 were processed using MS. The raw data were analyzed with Proteome Discoverer 2.1 and Xcalibur 4.6, and peptides were quantified by precursor ion area detection and normalized through peak integration of the proteotypic peptide [K].HGGPKDEER.[H]. Formylation levels were quantified in the following residues: (A) Thr55, (B) Thr59, (C) Arg70, (D) Thr89, (E) Lys92, (F) Arg116, (G) Thr136 and Lys137, and (H) Lys137. Data were expressed as Mean ± SD from at least three independent experiments. Two-way ANOVA was applied for statistical analysis with significance levels of * p < 0.05, ** p < 0.01, *** p < 0.001, **** p < 0.0001.

**Figure 7.**
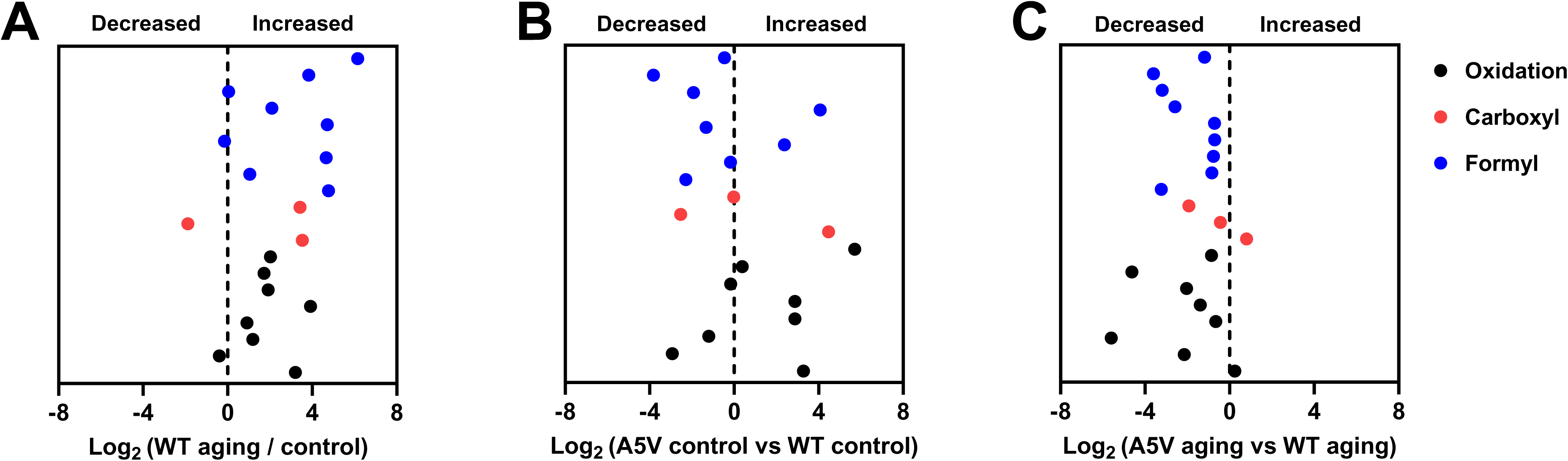
Aging increases damaging modifications in WT SOD1 and decreases in the A5V mutant. (A) Mean values of each damage modification in (A) aged WT SOD1 were compared to control WT SOD1 and plotted as log_2_ to express fold-change. (B) Mean values of each damage modification in control A5V SOD1 were compared to control WT SOD1 and plotted as log_2_ to express fold-change; (C) Mean values of each damage modification in aged A5V SOD1 were compared to aged WT SOD1 and plotted as log_2_ to express fold-change. Black dots indicate oxidative modifications, while red dots indicate carboxyl and black dots indicate formyl modifications.

### A5V mutation exhibited altered levels of damaging modifications than WT

The mutant SOD1 did not show alterations in the oxidation of the residues Trp33 (Figure 4a), His47 (Figure 4c), and His49 (Figure 4e) between the control and aging conditions. However, there was a decrease in the oxidation levels of Lys37 (Figure 4b), Asn87 (Figure 4g), and His111 (Figure 4h) after aging. A comparison between the control conditions of WT SOD1 and the A5V SOD1 mutant revealed a significant increase in oxidation at 4 out of 6 quantified residues (Trp33, Lys37, His47, and Asn87) of the mutant protein (Figure 4). This was accompanied by a decrease in His49, with no detectable change in His111. In contrast, the A5V mutant obtained from aged cells exhibited reduced oxidation levels, particularly in His residues, compared to WT SOD1 (Figure 4).

The quantification of carboxyl modification on A5V SOD1 revealed a significant increase in Lys92, with no alteration in Thr59 and Lys137 after aging compared to the control condition (Figure 5). Under control conditions, Thr59 carboxylation levels were observed to decrease in the SOD1 mutant relative to WT, whereas an increase was noted for Lys137 (Figure 5). After aging, the A5V SOD1 exhibited a reduction in carboxylation levels of Thr59 and Lys92 compared to WT SOD1 (Figure 5a-b).

The analysis of formylation on A5V SOD1, distinct from carboxylation and oxidation, has demonstrated a consistent pattern across most observed residues. Aging was associated with a significant increase in Thr59, Arg70, and Arg136 formylation in mutant SOD1, while no significant changes were noted in other residues. The mutant A5V, under control conditions, showed a decrease in formylation of Thr55, Thr59, and Lys92 compared to WT SOD1. Under aging conditions, the formylation levels of all observed residues in A5V SOD1 were significantly lower compared to WT SOD1 (Figure 6).

To evaluate whether there is a pattern of alteration in damaging PTMs, we conducted a qualitative analysis comparing different conditions. Interestingly, contrary to our expectations, our data revealed that after aging, A5V SOD1 exhibited less damage than WT SOD1 under the same conditions (Figure 7c). We attribute this phenomenon to A5V SOD1 inclusions, which were not concentrated during the purification process. These inclusions may sequester damaged SOD1 to prevent cellular apoptosis. The formation of aggregates of SOD1 mutants, mediated by aging, has been previously described by our group [24,30]. The comparison between A5V SOD1 and WT under control conditions does not indicate a definitive trend of increase or decrease in damaging PTMs; however, it does highlight a notable increase in oxidation, which may drive SOD1 aggregation (Figure 7b).

## Discussion

The SOD1 protein has been gaining more prominence when it comes to its relationship with the neurodegenerative disease ALS. Recently, a new drug known as Toffersen was approved in several countries (such as the United States and some European Union countries) for the treatment of ALS cases related to SOD1 mutations, based on the silencing of SOD1 phenotypes that present mutations [31,32], reinforcing the role that damage to this protein can lead to neurodegeneration. However, despite much talk about the relationship between ALS and SOD1 lesions, advances in this area have shown that oxidative damage in the WT SOD1 is also related to the development of ALS. Some studies with tissue samples from post-mortem patients with both forms of ALS (sporadic and familial) have shown the presence of inclusions of WT and mutant SOD1 [7]. Therefore, to better understand how this protein induces neurodegeneration and thus be able to develop more effective therapies, it is important to focus not only on the damage that SOD1 mutants cause to healthy cells, but also on which modifications in WT SOD1 in a stressful environment can lead to the development of ALS. Based on the discoveries from the last 5 years, the present work sought to analyze how oxidative stress induced by a chronological aging process can induce PTMs in WT SOD1 and A5V SOD1, an aggressive mutant, and how these modifications can influence ALS-related damage.

Both SOD1 forms, WT and mutant, were obtained from humanized yeast models, where the endogenous yeast SOD1 was replaced by the human ortholog. The yeast *S. cerevisiae* is a widely discussed model in ALS research, offering valuable insights into oxidative stress and its connection to the disease [33]. As an eukaryotic organism with a fully sequenced genome, yeast shares significant genetic similarities with humans; approximately one-third of yeast genes have a human ortholog, and two-thirds show homology with human genes [33]. Therefore, yeast proteins can replace their corresponding human proteins, like SOD1. Yeast models are also advantageous in studies investigating the effects of oxidative stress. This is because yeast metabolism can be adjusted by changing the carbon source in its growth medium. Yeast can switch between fermentative metabolism (which results in low oxidative stress) and respiratory metabolism (which results in high oxidative stress). This ability to modulate oxidative stress, by growing cells in stress-free conditions, makes yeast a valuable model for this type of study.

Initially, physiological changes, such as the phosphorylation levels of serine residues on SOD1, were evaluated (Figure 2b, c). Studies have shown that SOD1 phosphorylation on serine residues is important for SOD1 localization and protection against oxidative stress [26]. Phosphorylation on Ser60 and Ser99, after oxidative stress induction, signals a portion of cytoplasmic SOD1 to translocate to the nucleus, where it regulates the activation of antioxidant defense pathways [5]. Figure 2c shows that phosphorylation levels of WT SOD1 are higher than those of the mutant after cellular aging. Combining these phosphorylation results with literature findings on the nuclear localization levels of WT SOD1 and the A5V mutant—which indicate that nuclear levels of WT are higher than those of A5V [30] — it can be stated that higher contents of phosphorylated serine on SOD1 correspond to greater nuclear localization. In addition to phospho-Ser levels, the carbonylation levels of both SOD1 proteins were also measured. As shown in Figure 2b, the carbonylation levels of WT SOD1 increased after cellular aging and are statistically higher than those of the A5V mutant, which remained unchanged. While it may be surprising that WT SOD1 is more carbonylated than the mutant, this result corroborates previous findings about the oxidation level of WT SOD1 in models of sALS. According to Guareschi et al., samples of tissues from sALS patients show WT SOD1 in a hyperoxidized form, a condition that causes WT SOD1 to behave similarly to a SOD1 mutant [34]. Furthermore, studies have shown that, under the same conditions used here, A5V SOD1 forms inclusions, while WT does not [24]. This structural conformation may hinder access to the carbonylation sites of the A5V SOD1 mutant.

The literature has identified some PTMs in SOD1 using different models, which helps pinpoint the residues and types of PTMs that can occur in the protein. However, few studies compare the extent of these modifications in the protein. Moreover, research is lacking on how aging and ALS-related SOD1 mutations might alter the PTM profile of the protein. Using MS, it is possible to perform qualitative and quantitative analyses of the different PTMs present in the residues of WT and A5V SOD1. It has already been discussed in the literature that SOD1 can present more than 44 different conformations, depending on the variety and extent of PTMs, that can have physiological and pathological implications [13]. From the results obtained in this work, it was possible to identify physiological modifications (acetylation and succinylation) and damage (oxidation, formylation and carboxylation) in different residues distributed throughout the SOD1 sequence (Table 1 and S1).

Acetylation represents a prevalent PTM that is important to the regulation of various biological processes, including protein interactions, cell cycle progression, metabolic pathways, and the transport of proteins to the nucleus, alongside numerous other functions that have been extensively identified and characterized. Acetylation can occur in Lys, Ala, Arg, Asp, Cys, Gly, Glu, Met, Pro, Ser, Thr, and Val residues, but it is most commonly identified in lysine residues [35]. Studies have shown that acetylation in lysine residues is vital for cell development, and that dysregulation of this process can lead to the development of diseases, including neurodegenerative diseases such as Parkinson’s and Huntington’s [35].

Studies have shown that acetylation of the Lys71 residue of SOD1 decreases its enzymatic activity due to impaired interaction between CCS and SOD1. Additionally, acetylation of Lys123 inhibits SOD1’s ability to suppress cellular respiration, likely by disrupting the interaction between SOD1 and Ck1_γ_. This interaction may also be modulated by acetylation of Lys137 [5]. As shown in Figure 3, acetylation was identified in five Lys residues; however, as mentioned previously, only acetylation of Lys123 and Lys137 has a defined function related to the regulation of respiratory metabolism. While the Lys123 acetylation levels of WT SOD1 remained unchanged, a significant decrease was observed in its levels on A5V SOD1 after cellular aging. This result suggests that the SOD1 mutation in an oxidative environment contributes to the deacetylation of the residue, inducing a respiratory metabolism. A major source of intracellular ROS is the respiratory chain, and an uncontrolled increase in ROS can lead to increased cellular damage, including oxidative damage, which is a biomarker of aging and neurodegenerative diseases like ALS. This decrease in acetylation levels may also be related to the degree of A5V SOD1 aggregation. We can hypothesize that this relationship may be associated with two factors: a) Following aging, with the increase in the aggregation of mutant SOD1, the resulting protein structure may not be favorable for acetylation at these residues, reducing acetylation levels and promoting a respiratory metabolism, thus exacerbating ROS production and ultimately inducing intracellular oxidative stress; or b) The conformational change of SOD1 due to the presence of the A5V mutation, combined with metabolic changes in cells after cellular aging, induces deacetylation of the protein, increasing ROS concentration, which leads to intracellular oxidative damage, including contributing to the formation of mutant SOD1 inclusions. The remaining acetylation sites identified do not yet have defined functions for SOD1 in the literature, and future specific analyses of each residue would be ideal to understand how the studied conditions influence the variation in acetylation levels found in the present work and the function that SOD1 performs.

Another physiological modification detected was the succinylation of the Lys137 residue (Figure 3f). Succinylation, like acetylation, is a type of acylation that adds two carbon atoms to a lysine residue through a reaction with succinyl CoA, changing the surface charge from 0 to -1 [5]. Succinylation can occur in SOD1 at Lys10, Lys76, Lys92, Lys123, Lys129, and Lys137 (https://www.phosphosite.org/), but few of these have been explored in detail in the literature until now. Succinylation of Lys123 in SOD1 is thought to affect its enzymatic activity [5] and help regulate respiratory metabolism [29]. Although the observed differences between the conditions studied and between the two SOD1 isoforms make this an interesting finding, the function of succinylation at the Lys137 residue of SOD1 remains undefined in the literature. We propose that the significant increase in succinylation levels in wild-type SOD1 after aging, compared to the control, and the absence of change in A5V SOD1, suggests this PTM plays a role in cellular protection against oxidative stress, similar to the role of succinylation at Lys123.

While we could assess the levels of two physiological modifications by MS, most of the analyzed results pertain to potentially harmful PTMs. Modifications like oxidation, carbonylation, and formylation might be linked to oxidative stress caused by cellular aging in this study, particularly for WT SOD1. They may also be due to the A5V mutation in SOD1, which could lead to structural changes that make the protein more susceptible to damage under stress.

Cellular aging, inflammatory processes, epigenetic alterations, and proteostatic decline are characterized by several hallmarks, many of which are regulated by different PTMs, including oxidation [36]. Oxidation, also known as hydroxylation, is a marker of oxidative stress. Our work addresses a strategy for cellular aging that leads to increased oxidative stress, as demonstrated in previous studies by our group [24]. Oxidation of SOD1 residues can impair protein conformation, activity, and folding and is a predominant non-enzymatic PTM associated with aging [13]. The chemical changes resulting from oxidation can contribute to aggregation, a pathological hallmark observed in ALS patients [4]. Our results identified several oxidized residues along the SOD1 sequence, including Trp33, Lys37, His47, His49, His81, His111, Asn87, N132, and N140 (Figure 4). Oxidation at tryptophan residues, specifically at residue 33, promotes the formation of ditryptophan covalent bonds between SOD1 monomers. This results in unfolding, oligomerization, and non-amyloid aggregation of SOD1 [37]. Our results demonstrate a significant increase in oxidation at the Trp33 residue of WT SOD1 during stress conditions (Figure 4a). This increase may correlate with the development of WT SOD1 aggregates found in sALS patients [7]. Furthermore, oxidation levels in the A5V mutant remained constant after cellular aging but were higher than those observed in the control condition of WT SOD1. This suggests that the A5V mutation increases susceptibility to oxidation at this residue compared to the wild-type protein, although not higher than the levels seen in WT SOD1 after induction of oxidative stress.

Histidines, more specifically His47, His49, His64, His72, His81, and His121, are residues that bind to the metal ions Zn^2+^ and Cu^2+^ present in SOD1, which play a structural and catalytic role, respectively. Damage to these residues can result in the breakdown of the residue-ion interaction, resulting in the loss of protein stability and conformation, increasing the propensity for the formation of SOD1 aggregates [5, 38]. Increased oxidation at His47, His49, His81, His111 and His121 residues observed for WT SOD1 (Figure 4) can be correlated with the development of SOD1 aggregates in sALS patients, resulting from the loss of protein stability, from the chemical modification in the side chains that shift the conformational stability to the aggregation-prone state [13]. When comparing the levels for the mutant, no significant increases in oxidation were observed in these residues under the different conditions, which may reflect that the loss of stability and propensity for aggregation of the A5V SOD1 resulted from a loss of intrinsic conformation of the protein mutation.

Damage such as carboxylation and formylation, present in the protein side chain, can also affect stability and conformation, from the addition of CO_2_ and CO groups, respectively. Carboxylation of lysine residues occurs through the interaction of CO_2_ or CO_3_ groups with the amine group of the residue, influenced by the environment and the structural conformation of the protein [39]. Formylation is a PTM similar to acetylation, but with a smaller mass addition (28 Da, while acetylation is 42 Da). In some cases, the formyl group may appear as a consequence of the treatment of the samples with formic acid and is considered an artifact and not a PTM [40]. However, since our samples were treated with trifluoroacetic acid, and not with formic acid, we consider the observed formylation as a PTM. As seen in Figures 5 and 6, respectively, in general, carboxylation and formylation damage showed a significant increase during oxidative stress for most of the identified WT SOD1 residues, showing an increase in oxidative damage that may justify the role of WT SOD1 in the sporadic form of ALS. Among the residues that showed increases in these damages after aging (Thr55, Arg70, Thr89, Lys92, Arg116, Lys129, Thr136 and Lys137), it is possible to visualize a distribution between regions of the segment that comprises the broad Zn loop and the electrostatic loop of the SOD1 structure, regions responsible for the stabilization of the SOD1 structure [5]. The loss of structural stability may induce a decrease in enzymatic activity and increase the probability of formation of damaged WT SOD1 inclusions. Unlike what was observed in WT SOD1, the carboxylation and formylation levels of A5V SOD1 remained the same in control and aging conditions in most of the results, suggesting that the A5V mutation is already sufficient for SOD1 to suffer damage, even under normal conditions.

Most of the results obtained for A5V SOD1 did not show any variation in the levels of modifications under control and aging conditions, especially those related to oxidative damage. Although an increase in damage is expected from SOD1 that already has a toxic mutation, the maintenance of PTM levels may be related to the degree of agglomeration. Studies already published by our group show that the A5V mutant, when subjected to chronological aging conditions similar to those used in this study, presents an increase in total aggregation, when compared to the control condition [24]. SOD1 inclusions are formed by unfolded and poorly structured monomers, so that they are unable to maintain the stabilized dimer structure, and begin to interact in a disorganized manner, forming trimers, oligomers, and inclusions [41]. As previously discussed, oxidative damage, carboxylation, and formylation, for example, contribute to the misfolding of SOD1, which leads to the formation of inclusions. Thus, it is expected that most of the A5V SOD1 that suffered damage due to aging is in the form of inclusions, and not in its free dimeric form. However, the SOD1 enrichment technique used in this work could not capture SOD1 aggregates, so that the samples analyzed by both western blotting and MS experiments only considered A5V SOD1 in the dimeric form, and therefore, most of the levels observed for the mutant protein did not vary, even after cellular aging. In contrast, the values observed for WT SOD1 varied when comparing the control and aging conditions. The aging condition used in this work is not sufficient to induce an aggregation in the wild-type form of SOD1 [24], so that during the enrichment technique, a good part of the SOD1 is recovered, ensuring that the results obtained and discussed here reflect as closely as possible the reality with precision and certainty what occurs in the control and aging conditions for WT SOD1.

It is interesting to observe the values of oxidation, carbonylation, and formylation levels of WT SOD1 after aging and A5V SOD1 control. In most results, the levels remained very close or higher for WT SOD1 (Figure 7). These results indicate that despite not inducing the formation of WT SOD1 aggregates, the treatment used was sufficient to induce an increase in the oxidation levels of WT SOD1 comparable to the oxidation degree of mutant SOD1 in the control condition. Our results confirm what some studies have shown: WT SOD1 in sALS models is in the hyperoxidized form compared to mutant SOD1 [34]. These modifications in the oxidation state of WT SOD1 help to understand why even a SOD1 without any type of mutation in its structure and maintaining enzymatic activity, contributes to the development and propagation of ALS, and why in some more extreme cases of stress, it can be found forming aggregates [7,24].

In summary, our data identify and quantify, for the first time to our knowledge, the types and levels of PTMs of WT SOD1 in response to aging, a risk factor for sALS. This work also investigated the effect of the severe A5V mutation on SOD1 PTMs. Our results support previous findings in the literature regarding the oxidation levels of WT SOD1 and its connection to sALS. We found that certain physiological PTMs, like acetylation and succinylation, which currently have no known role in SOD1, show different levels depending on both the presence of mutations in the protein and the environment around SOD1. More research to understand the importance of these PTMs could help us better understand SOD1’s role in healthy states and how their absence might contribute to ALS development.

## Material and Methods

### Yeast strains and Growth Conditions

Knockout *SOD1* yeast cells (*MATa; his3, leu2, met15; ura3; sod1::KanM*X4), acquired from Euroscarf were transformed with YEp plasmids containing cDNA sequences of human WT and A5V SOD1, as previously described [16]. Cells were grown at 28 °C and 160 rpm until they reached the mid-exponential phase in a dropout medium containing 2% glucose but lacking leucine (control condition). For the chronological aging condition, which measures the lifespan of non-dividing yeast populations, cells were harvested by centrifugation from the culture medium, washed twice with sterile water, resuspended in the same volume of water, and incubated at 37 °C and 160 rpm for 24 hours [23].

### Purification of hSOD1

Soluble proteins from crude extract were obtained by treating the cells with lysis buffer (200 mM Tris-HCl, 0.1 mM EDTA, 50 mM NaCl, pH 8.0), protease inhibitor mixture (cOmplete, mini, EDTA-free protease mixture inhibitor cocktail, Sigma-Aldrich, Waltham, MA, USA) and submitted to 4 cycles of 1 minute on vortex and 1 minute on ice. After obtaining the extract, protein precipitation was performed using ammonium sulfate ((NH_4_)_2_SO_4_) for 20 minutes in an ice bath, the solution was centrifuged at 15000 xg at 4 °C, and the supernatant collected. 0.25 volume of saturated (NH_4_)_2_SO_4_ solution at 25 °C was added to the supernatant to adjust the solution saturation to 64 % and the solution was centrifuged again at 15000 xg at 4 °C and the supernatant collected. To remove excess salt in solution, it was performed dialysis for 48 h in 50 mM Tris-HCl pH 7 buffer, using a 25 mm cellulose membrane (Merck - D9777-100FT). The supernatant was concentrated with Amicon Ultra-30 kDa-Merck Millipore. Ion exchange chromatography was performed using a HiPrep™ DEAE Fast Flow 16/10 anion column operating at a flow rate of 0.5 ml/min, collecting the first fraction after the start of elution using a 1 M KCl gradient. Fractions containing SOD1 were confirmed by activity analysis. Finally, the sample was concentrated using Amicon Ultra-10 kDa-Merck Millipore. Protein concentration was determined by the Bradford assay [23].

### Superoxide dismutase activity

Protein samples were applied in 15 % native polyacrylamide gels, which were run at 200 V for 2 h. SOD1 activity was determined using the nitroblue tetrazolium (CHEM-IMPEX INT’L INC. Lot# 001690-2017041001) method, as previously described [24]. For SOD1 enzyme activity quantification, the digital images were analyzed by Fiji ImageJ software.

### Immunoblotting

Protein samples were fractionated by SDS/PAGE, using 15 % polyacrylamide gels and transferred to a nitrocellulose membrane (0.45 µm, 300 mm x 4 m - GE healthcare). The blots were incubated overnight in 3 % BSA solution at room temperature, then incubated with primary antibody for 1 h. The antibodies used were: anti-SOD1 (1:2000 - Sigma-Aldrich, produced in rabbit - HPA001401), anti-phosphoserine (1:1000 - Sigma-Aldrich, produced in rabbit - SAB5200086) and anti-DNP (1:1000, Sigma-Aldrich Merck, produced in rabitt - D9656). To determine the levels of carbonyl, which are detected with anti-DNP, the blots were previously treated with 15 mL of dinitrophenylhydrazine (DNPH) 2 mM solution in 2 M HCl, for 15 min, at room temperature, then washed once with 5 M HCl and twice with PBS. Next, the membranes were washed twice to remove the primary antibody and then incubated for 1 h, with anti-rabbit secondary antibodies conjugated to horseradish peroxidase. To determine the levels of phospho-serine and carbonyl on SOD1, after mark with anti-SOD1 and secondary antibody, each membrane was stripped using a stripping buffer (1.5% glycine, 0.1% SDS and 1% tween, pH 2.2). The process was performed 2x with 10 mL of buffer for 15 minutes followed by three washes with PBST. After stripping, the membrane was blocked again with PBST-BSA 3% and then labeled with the next antibody of interest. For detection of the bands, the blots were treated with luminol reagent and peroxide solution kit (Western blotting ECL-Promega) and then scanned in Fusion Solo 6S imaging system (Vilber Lourmat, Marne-la-Vallee, France). The band intensity was quantified using the Fiji ImageJ software.

### Sample processing for mass spectrometry analysis

The experimental layout used to identify and quantify PTMs observed in WT and A5V human SOD1 in response to aging is shown in Figure 1. The purified protein extract was precipitated overnight with ice-cold acetone (100 %) at a 4:1 ratio (acetone/sample, v/v). After precipitation, the solution was centrifuged at 20,000 x g at 4 °C for 30 min, and the resulting pellet was washed twice with ice-cold acetone, centrifuging under the same conditions. The pellet was then completely dried in a SpeedVac vacuum concentrator (Savant) and dissolved in a solution containing 7 M urea, 2 M thiourea, and 200 mM triethylammonium bicarbonate (TEAB). Protein concentration was determined using a Qubit 4.0® fluorometer (Invitrogen – Thermo Fisher Scientific - Q33238), following the manufacturer’s instructions. For protein digestion, 50 µg of protein was reduced with 10 mM dithiothreitol for 1 h at 30 °C and alkylated with 40 mM iodoacetamide for 30 min in the dark at room temperature. Subsequently, the samples were diluted 10-fold in 500 mM TEAB and incubated with trypsin (Promega, Madison, WI USA – V511A) at a trypsin-to-protein ratio of 1:50 for 16 h at 37 °C. Digestion was stopped by adding trifluoroacetic acid (TFA) to a final concentration of 10 %. Samples were then cleaned using reversed-phase microcolumns with POROS ™ 20 R2 resin (Thermo Scientific™) and C18 disk (Empore™ 3M). Peptides were eluted with 0.1 % TFA in 50 % acetonitrile (ACN), followed by 0.1% TFA in 70% ACN. The peptides were completely dried in a SpeedVac and resuspended in 0.1 % formic acid solution for subsequent mass spectrometry (MS) analysis.

### Determination of PTMs by MS and Data Analysis

LC-MS/MS analyses were performed using an Easy-nLC 1000 system coupled to a Q-Exactive Plus mass spectrometer (Thermo Scientific, Waltham, MA, USA). Ionization was conducted using an electrospray ion source in positive polarity under data-dependent acquisition (DDA) mode, with a spray voltage of 2.5 kV and a capillary temperature of 200 °C. The spectrometer was set to acquire Full MS spectra in the range of 350 to 2000 m/z, with a resolution of 70,000 (m/z 200) and an automatic gain control (AGC) target of 1e6. The 15 most intense ions were selected for fragmentation in dd-MS² mode with a normalized collision energy (NCE) of 30, a resolution of 17,500, an isolation window of 2.0 m/z, and a dynamic exclusion of 45 seconds. The spectra generated from LC-MS/MS analyses were processed using Proteome Discoverer 2.1 software (Thermo Scientific) with the Sequest HT search engine, using the protein database (UniProt). The following parameters were applied: precursor mass tolerance of 10 ppm, fragment mass tolerance of 0.1 Da, trypsin specificity with up to two missed cleavages allowed. Fixed modifications included carbamidomethylation, while variable modifications included oxidation, acetylation, succinylation, phosphorylation, carboxylation, and ubiquitination. High-confidence peptides were selected, and only identifications with q values ≤ 0.01 (FDR ≤ 1%) were considered. FDR values were calculated using the Target Decoy PSM strategy.

The intensity of the area of the modified peptides was obtained using the Xcalibur^TM^ 4.3 software (Thermo Scientific). The areas of the peptides of interest were integrated into the chromatograms, filtering by the retention time and mass/charge ratio of each peptide analyzed. The data were normalized with the area of the non-modified prototypic peptide of SOD1.

### Bioinformatic analysis

Data was quantified in software Perseus version 2.0.7.0 [25]. The normalized data from all biological triplicates were transformed to Log2. The values obtained with the normalization of the WT SOD1 and A5V SOD1 triplicates were compared with each other using Pearson’s correlation. To detect the main effect of normalization on the data, a principal component analysis (PCA) was performed for the normalized data and raw data.

### Statistical analysis

Data were analyzed using GraphPad Prism 8 software and expressed as Mean ± SD of at least 3 independent experiments. Statistical differences were calculated using two-way ANOVA, as described in figure subtitle. Significance was expressed: \**P* < 0.05, \*\**P* < 0.01, \*\*\**P* < 0.001 and **** *P* < 0.0001.

## Authors Contributions

G.F.S: Data collection, analysis, Writing - original draft; R.S.S.M; Conceptualization, analysis, Writing - original draft; J.R.M.N: Analysis, Writing - review & editing; M.M.: Data collection, analysis; C.F.: Resources, supervision, funding acquisition; M.J.: Resources, supervision, funding acquisition; E.C.A.E: Conceptualization, Resources, Writing – review & editing, Supervision, Project administration, Funding acquisition.

## References

[1] López-Otín C & Kroemer G (2021) Hallmarks of health. Cell 184(1), p. 33–63, doi:10.1016/j.cell.2020.11.034.

[2] Ebert T, Tran N, Schurgers L, Stenvinkel P & e Shiels PG (2022) Ageing - Oxidative Stress, PTMs and Disease. Molecular Aspects of Medicine 86 (101099), 101099, doi:10.1016/j.mam.2022.101099.

[3] Kang W, Liu H, Hu Q, Wang L, Liu J, Zheng Z, Zhang W, Ren J, Zhu F & Liu GH (2022) Epigenetic regulation of aging: implications for interventions of aging and diseases. Signal transduction and targeted therapy 7(1), p. 374, doi:10.1038/s41392-022-01211-8.

[4] Hardiman O, Al-Chalabi A, Chio A, Corr EM, Logroscino G, Robberecht W, Shaw PJ, Simmons Z & Van den Berg LH (2017) Amyotrophic lateral sclerosis. Nat Rev Dis Primers 3, 17071, doi:10.1038/nrdp.2017.71.

[5] Eleutherio EC A, Magalhães RSS, Brasil AA, Monteiro Neto JR & Paranhos LH (2021) SOD1, more than just an antioxidant. Archives of Biochemistry and Biophysics 697(108701), 108701, doi:10.1016/j.abb.2020.108701.

[6] Proctor EA, Ding F & Dokholyan NV (2011) Structural and thermodynamic effects of post-translational modifications in mutant and wild type Cu, Zn superoxide dismutase. Journal of Molecular Biology 408(3), 555–567, doi:10.1016/j.jmb.2011.03.004.

[7] Trist BG, Genoud S, Roudeau S, Rookyard A, Abdeen A, Cottam V, … Double KL (2022). Altered SOD1 maturation and post-translational modification in amyotrophic lateral sclerosis spinal cord. Brain: A Journal of Neurology 145(9), 3108–3130, doi:10.1093/brain/awac165.

[8] Kalia M, Miotto M, Ness D, Opie-Martin S, Spargo TP, Di Rienzo L, … Iacoangeli A (2023) Molecular dynamics analysis of superoxide dismutase 1 mutations suggests decoupling between mechanisms underlying ALS onset and progression. Computational and Structural Biotechnology Journal 21, 5296–5308, doi:10.1016/j.csbj.2023.09.016.

[9] Valentine JS, Doucette PA & Zittin Potter S (2005) Copper-zinc superoxide dismutase and amyotrophic lateral sclerosis. Annual Review of Biochemistry 74(1), 563–593, doi:10.1146/annurev.biochem.72.121801.161647.

[10] Kaur SJ, McKeown SR & Rashid S (2016) Mutant SOD1 mediated pathogenesis of Amyotrophic Lateral Sclerosis. Gene 577(2), 109–118, doi:10.1016/j.gene.2015.11.049.

[11] Lin C, Zeng H, Lu J, Xie Z, Sun W, Luo C, Ding J, Yuan S, Geng M & Huang M (2015) Acetylation at lysine 71 inactivates superoxide dismutase 1 and sensitizes cancer cells to genotoxic agents. Oncotarget 6[24], 20578–20591, doi:10.18632/oncotarget.3987.

[12] Lin ZF, Xu HB, Wang JY, Lin Q, Ruan Z, Liu FB, Jin W, Huang HH & Chen X (2013) SIRT5 desuccinylates and activates SOD1 to eliminate ROS. Biochemical and Biophysical Research Communications, 441(1), 191–195, doi:10.1016/j.bbrc.2013.10.033.

[13] Shafiq K, Sanghai N, Guo Y & Kong J (2021) Implication of post-translationally modified SOD1 in pathological aging. GeroScience 43(2), 507–515, doi:10.1007/s11357-021-00332-2.

[14] Banks CJ & Andersen JL (2019) Mechanisms of SOD1 regulation by post-translational modifications. Redox Biology, 26(101270), 101270, doi:10.1016/j.redox.2019.101270.

[15] Tsang CK, Chen M, Cheng X, Qi Y, Chen Y, Das I, Li X, Vallat B, Fu LW, Qian CN, Wang HY, White E, Burley SK & Zheng XFS (2018) SOD1 phosphorylation by mTORC1 couples nutrient sensing and redox regulation. Molecular Cell 70(3), 502–515.e8, doi:10.1016/j.molcel.2018.03.029.

[16] Paranhos LH, Magalhães RSS, Brasil AA, Monteiro Neto JR, Ribeiro GD, Queiroz DD, Santos VM, & Eleutherio ECA (2024) The familial amyotrophic lateral sclerosis-associated A4V SOD1 mutant is not able to regulate aerobic glycolysis. Biochimica et Biophysica Acta. General Subjects 1868(8), 130634, doi:/10.1016/j.bbagen.2024.130634.

[17] Percio A, Cicchinelli M, Masci D, Summo M, Urbani A & Greco V (2024) Oxidative cysteine post translational modifications drive the redox code underlying neurodegeneration and amyotrophic lateral sclerosis. Antioxidants (Basel, Switzerland) 13(8), 883, doi:10.3390/antiox13080883.

[18] Bedja-Iacona L, Richard E, Marouillat S, Brulard C, Alouane T, Beltran S, Andres CR, Blasco H, Corcia P, Veyrat-Durebex C & Vourc’h P (2024) Post-translational variants of major proteins in amyotrophic lateral sclerosis provide new insights into the pathophysiology of the disease. International Journal of Molecular Sciences 25(16), 8664, doi:10.3390/ijms25168664.

[19] Dunphy K, Dowling P, Bazou D & O’Gorman (2021) Current methods of post-translational modification analysis and their applications in blood cancers. Cancers 13(8), 1930, doi:10.3390/cancers13081930.

[20] Azevedo R, Jacquemin C, Villain N, Fenaille F, Lamari F & Becher F (2022) Mass spectrometry for neurobiomarker discovery: The relevance of post-translational modifications. Cells (Basel, Switzerland) 11(8), 1279, doi:10.3390/cells11081279.

[21] Chen C, Hou J, Tanner JJ & Cheng J (2020) Bioinformatics methods for mass spectrometry-based proteomics data analysis. International Journal of Molecular Sciences 21(8), 2873, doi:10.3390/ijms21082873.

[22] Giannakou M, Akrani I, Tsoka A, Myrianthopoulos V, Mikros E, Vorgias C & Hatzinikolaou DG (2024) Discovery of novel inhibitors against ALS-related SOD1(A4V) aggregation through the screening of a chemical library using differential scanning fluorimetry (DSF). Pharmaceuticals (Basel, Switzerland) 17(10), 1286, doi:10.3390/ph17101286.

[23] Monteiro Neto JR, Ribeiro GD, Magalhães RSS, Follmer C, Outeiro TF & Eleutherio ECA (2023) Glycation modulates superoxide dismutase 1 aggregation and toxicity in models of sporadic amyotrophic lateral sclerosis. Biochimica et Biophysica Acta. Molecular Basis of Disease 1869(8), 166835, doi:10.1016/j.bbadis.2023.166835.

[24] Magalhães RSS, Monteiro Neto JR, Ribeiro GD, Paranhos LH & Eleutherio ECA (2024) Trehalose protects against superoxide dismutase 1 proteinopathy in an amyotrophic lateral sclerosis model. Antioxidants (Basel, Switzerland) 13(7), 807, doi:10.3390/antiox13070807.

[25] Tyanova S, Temu T, Sinitcyn P, Carlson A, Hein MY, Geiger T, Mann M & Cox J (2016) The Perseus computational platform for comprehensive analysis of (prote)omics data. Nature Methods 13(9), 731–740, doi:10.1038/nmeth.3901.

[26] Tsang CK, Liu Y, Thomas J, Zhang Y & Zheng XFS (2014) Superoxide dismutase 1 acts as a nuclear transcription factor to regulate oxidative stress resistance. Nature Communications 5(1), 3446, doi:10.1038/ncomms4446.

[27] Brasil AA, De Carvalho MDC, Gerhardt E, Queiroz DD, Pereira MD, Outeiro TF & Eleutherio ECA (2019) Characterization of the activity, aggregation, and toxicity of heterodimers of WT and ALS-associated mutant Sod1. Proceedings of the National Academy of Sciences of the United States of America 116(51), 25991–26000, doi:10.1073/pnas.1902483116.

[28] Weinert BT, Schölz C, Wagner SA, Iesmantavicius V, Su D, Daniel JA & Choudhary C (2013) Lysine succinylation is a frequently occurring modification in prokaryotes and eukaryotes and extensively overlaps with acetylation. Cell Reports 4(4), 842–851, doi:10.1016/j.celrep.2013.07.024.

[29] Banks CJ, Rodriguez NW, Gashler KR, Pandya RR, Mortenson JB, Whited MD, Soderblom EJ, Thompson JW, Moseley MA, Reddi AR, Tessem JS, Torres MP, Bikman BT & Andersen JL (2017) Acylation of superoxide dismutase 1 (SOD1) at K122 governs SOD1-mediated inhibition of mitochondrial respiration. Molecular and Cellular Biology 37[20], doi:10.1128/MCB.00354-17.

[30] Brasil AA, Magalhães RSS, De Carvalho MDC, Paiva I, Gerhardt E, Pereira MD, Outeiro TF & Eleutherio ECA (2018) Implications of fALS mutations on Sod1 function and oligomerization in cell models. Molecular Neurobiology 55(6), 5269–5281, doi:10.1007/s12035-017-0755-4.

[31] Miller TM, Cudkowicz ME, Genge A, Shaw PJ, Sobue G, Bucelli RC, … VALOR and OLE Working Group (2022) Trial of antisense oligonucleotide tofersen for SOD1 ALS. The New England Journal of Medicine 387(12), 1099–1110, doi:10.1056/NEJMoa2204705.

[32] Meyer T, Schumann P, Weydt P, Petri S, Koc Y, Spittel S, Bernsen S, Günther R, Weishaupt JH, Dreger M, Kolzarek F, Kettemann D, Norden J, Boentert M, Vidovic M, Meisel C, Münch C, Maier A & Körtvélyessy P (2023) Neurofilament light-chain response during therapy with antisense oligonucleotide tofersen in SOD1-related ALS: Treatment experience in clinical practice. Muscle & Nerve 67(6), 515–521, doi:10.1002/mus.27818.

[33] Di Gregorio SE & Duennwald ML (2018) ALS yeast models-past success stories and new opportunities. Frontiers in Molecular Neuroscience 11, 394, doi:10.3389/fnmol.2018.00394.

[34] Guareschi S, Cova E, Cereda C, Ceroni M, Donetti E, Bosco DA, Trotti D & Pasinelli P (2012) An over-oxidized form of superoxide dismutase found in sporadic amyotrophic lateral sclerosis with bulbar onset shares a toxic mechanism with mutant SOD1. Proceedings of the National Academy of Sciences of the United States of America 109(13), 5074–5079, doi:10.1073/pnas.1115402109.

[35] Ramazi S & Zahiri J (2021) Posttranslational modifications in proteins: resources, tools and prediction methods. Database: The Journal of Biological Databases and Curation 2021, doi:10.1093/database/baab012.

[36] Baldensperger T, Eggen M, Kappen J, Winterhalter PR, Pfirrmann T & Glomb MA (2020) Comprehensive analysis of posttranslational protein modifications in aging of subcellular compartments. Scientific Reports 10(1), 7596, doi:10.1038/s41598-020-64265-0.

[37] Coelho FR, Iqbal A, Linares E, Silva DF, Lima FS, Cuccovia IM & Augusto O (2014) Oxidation of the tryptophan 32 residue of human superoxide dismutase 1 caused by its bicarbonate-dependent peroxidase activity triggers the non-amyloid aggregation of the enzyme. The Journal of Biological Chemistry 289(44), 30690–30701, doi:10.1074/jbc.M114.586370.

[38] Ashkaran F, Seyedalipour B, Baziyar P & Hosseinkhani S (2024) Mutation/metal deficiency in the “electrostatic loop” enhanced aggregation process in apo/holo SOD1 variants: implications for ALS diseases. BMC Chemistry 18(1), 177, doi:10.1186/s13065-024-01289-x.

[39] Cain JA, Solis N & Cordwell SJ (2014) Beyond gene expression: the impact of protein post-translational modifications in bacteria. Journal of Proteomics 97, 265–286, doi:10.1016/j.jprot.2013.08.012.

[40] Alzate, O. Neuroproteomics. Londres, England: CRC Press, 2010, ISBN-13: 978-1-4200-7625-7.

[41] Choi ES & Dokholyan NV (2021) SOD1 oligomers in amyotrophic lateral sclerosis. Current Opinion in Structural Biology 66, 225–230, doi:10.1016/j.sbi.2020.12.002.

