## Supplementary material for "The effect of aging on post-translational modifications of wild-type human SOD1 and the A5V ALS mutant"

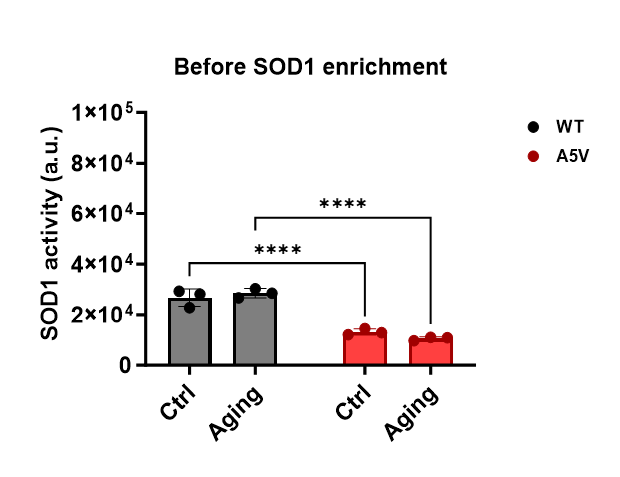

**Figure S1**. SOD1 activity of cell extracts obtained from cells cultured under fermentative conditions, with low levels of ROS (Ctrl), and after chronological aging (Ag), which increases oxidative stress.

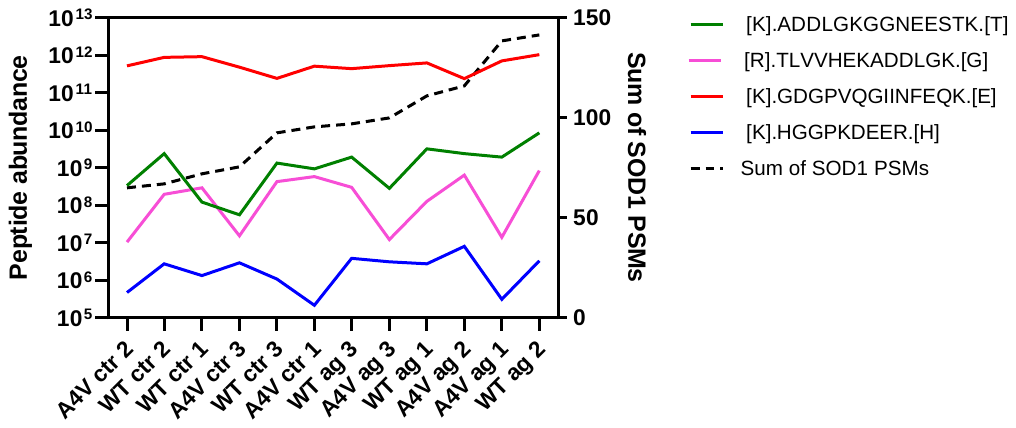

**Figure S2.** Relationship between the abundance of the proposed normalizing peptides with the sum of the total peptide spectrum matches (PSMs) of SOD1 in the samples.

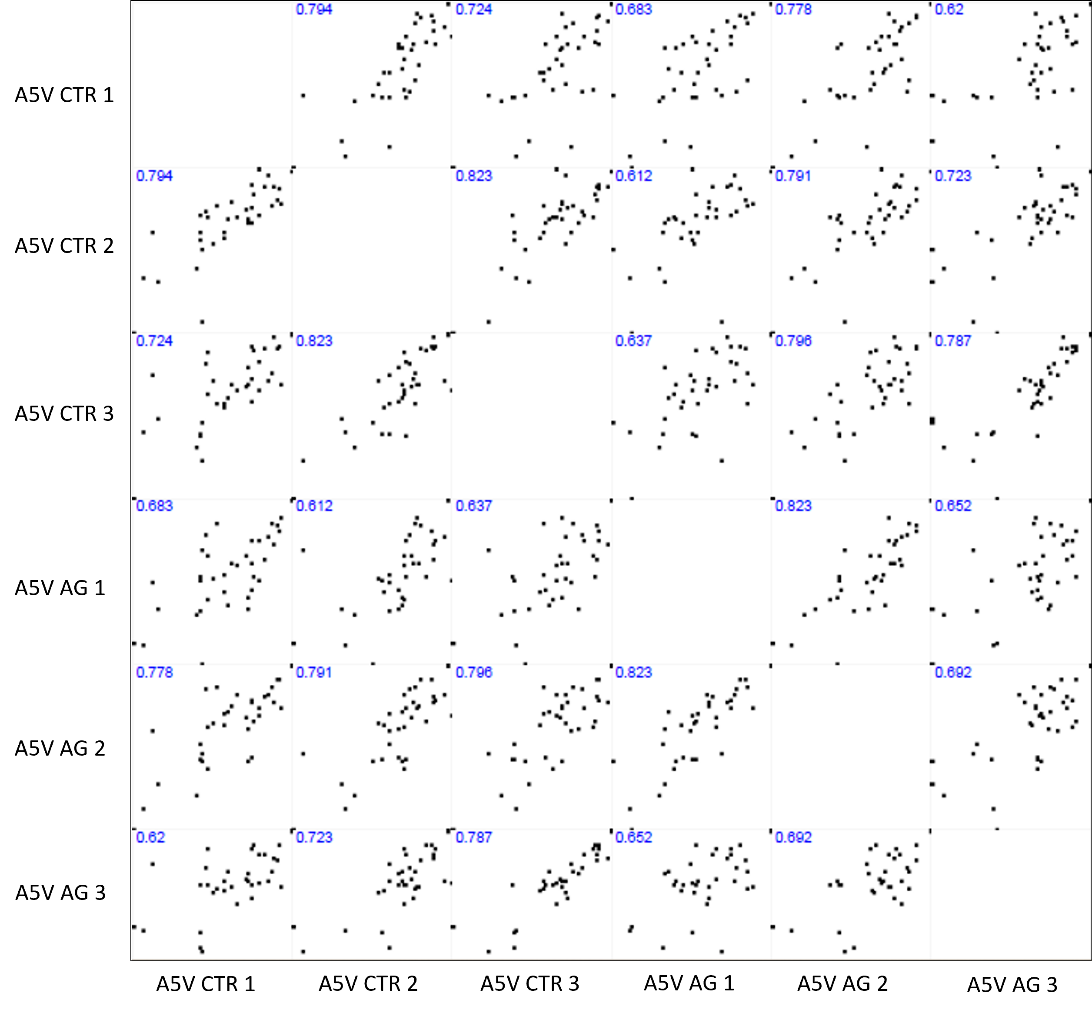

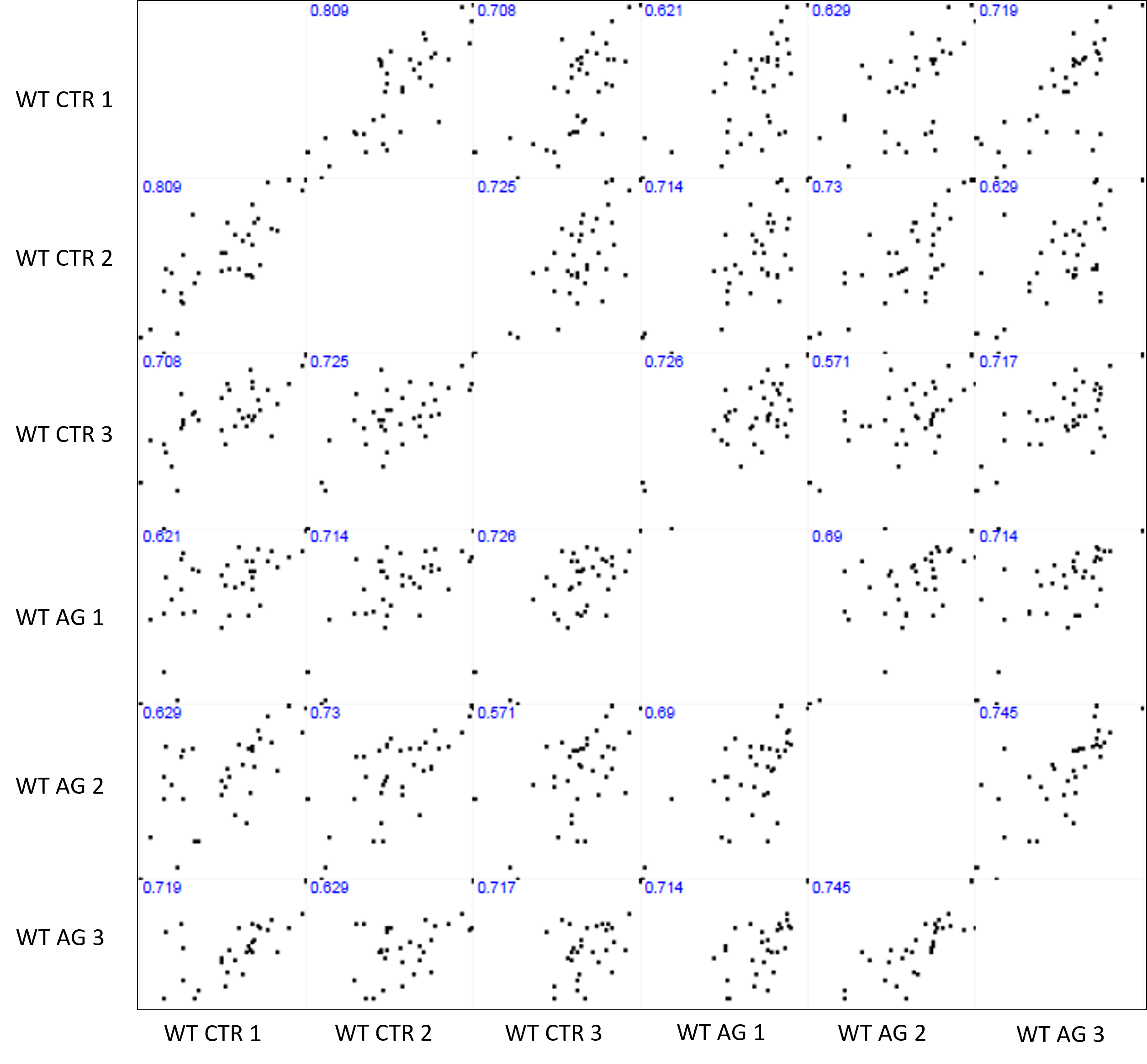

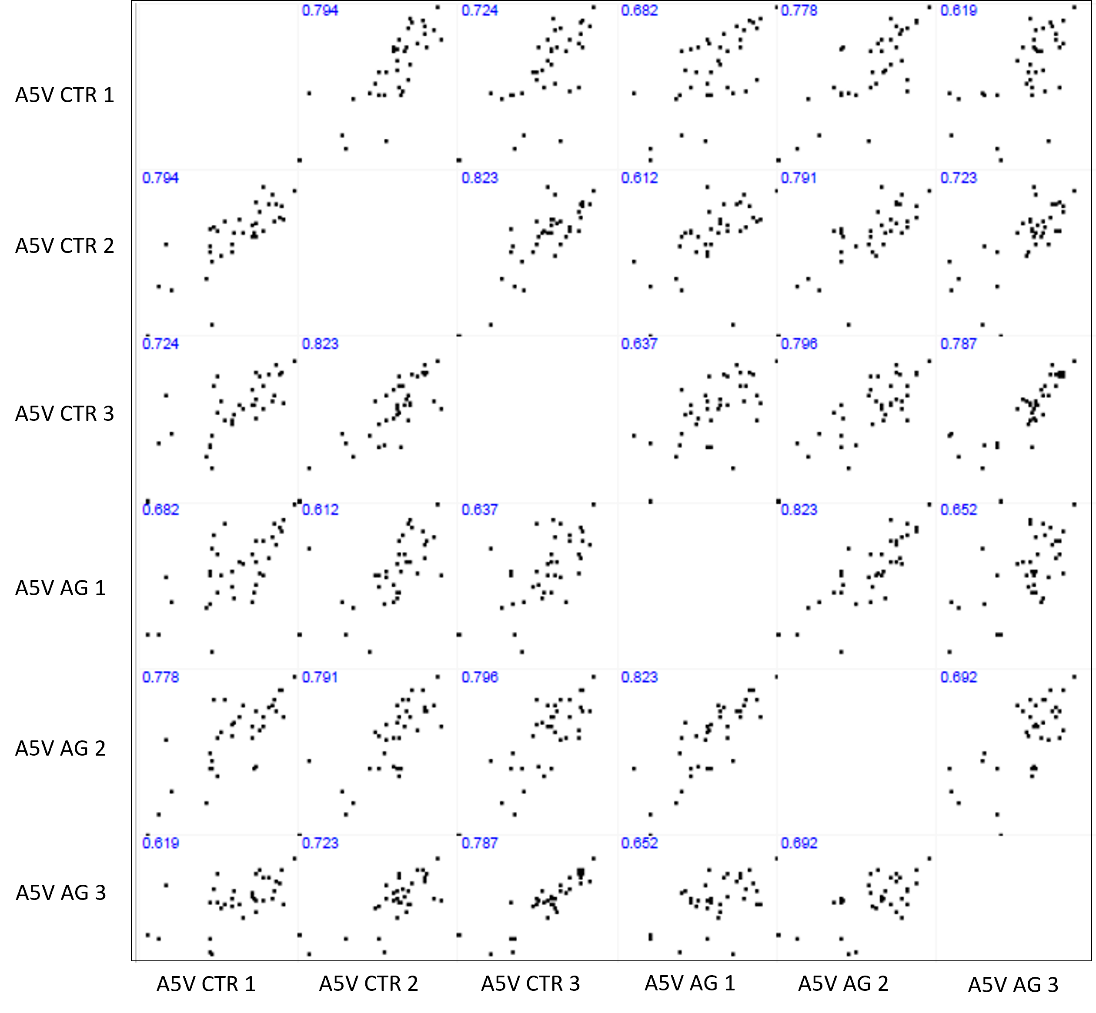

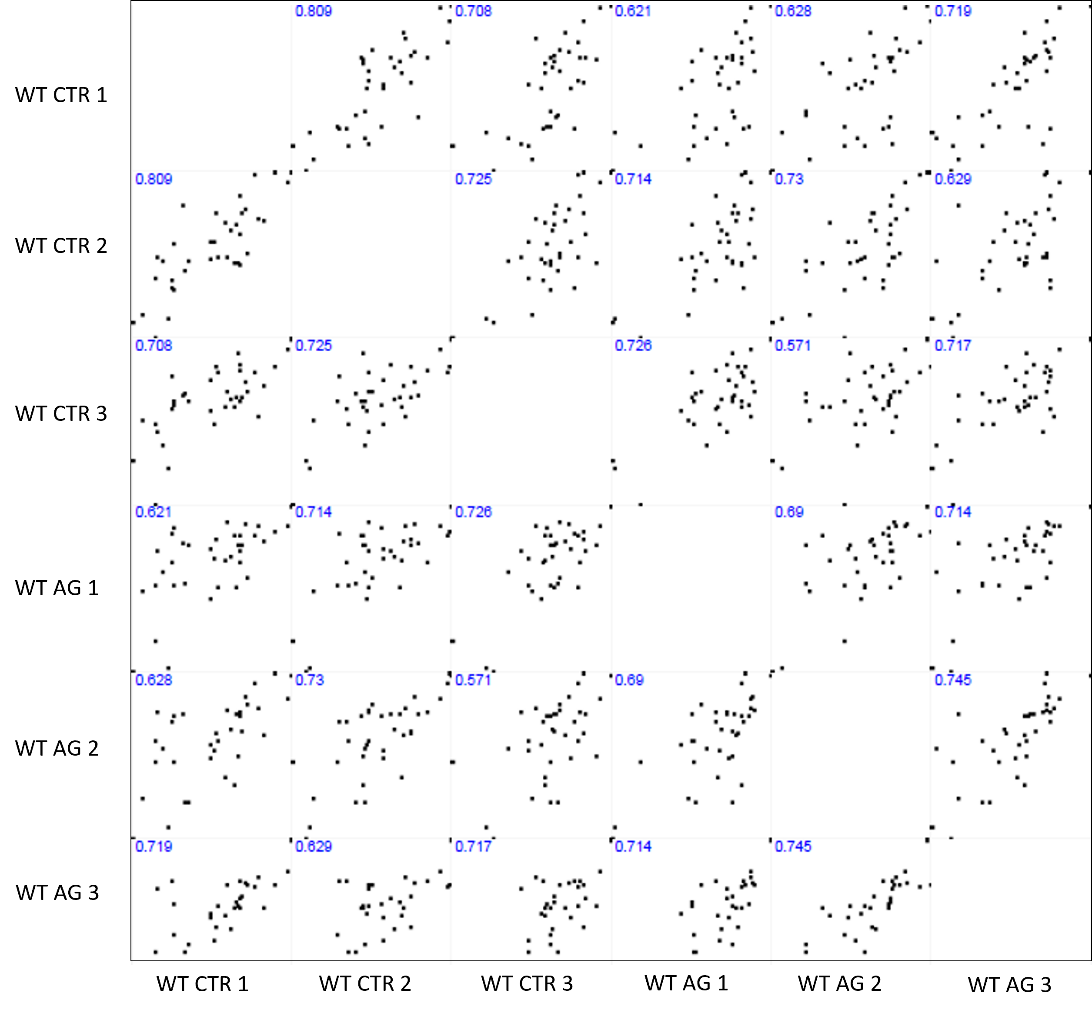

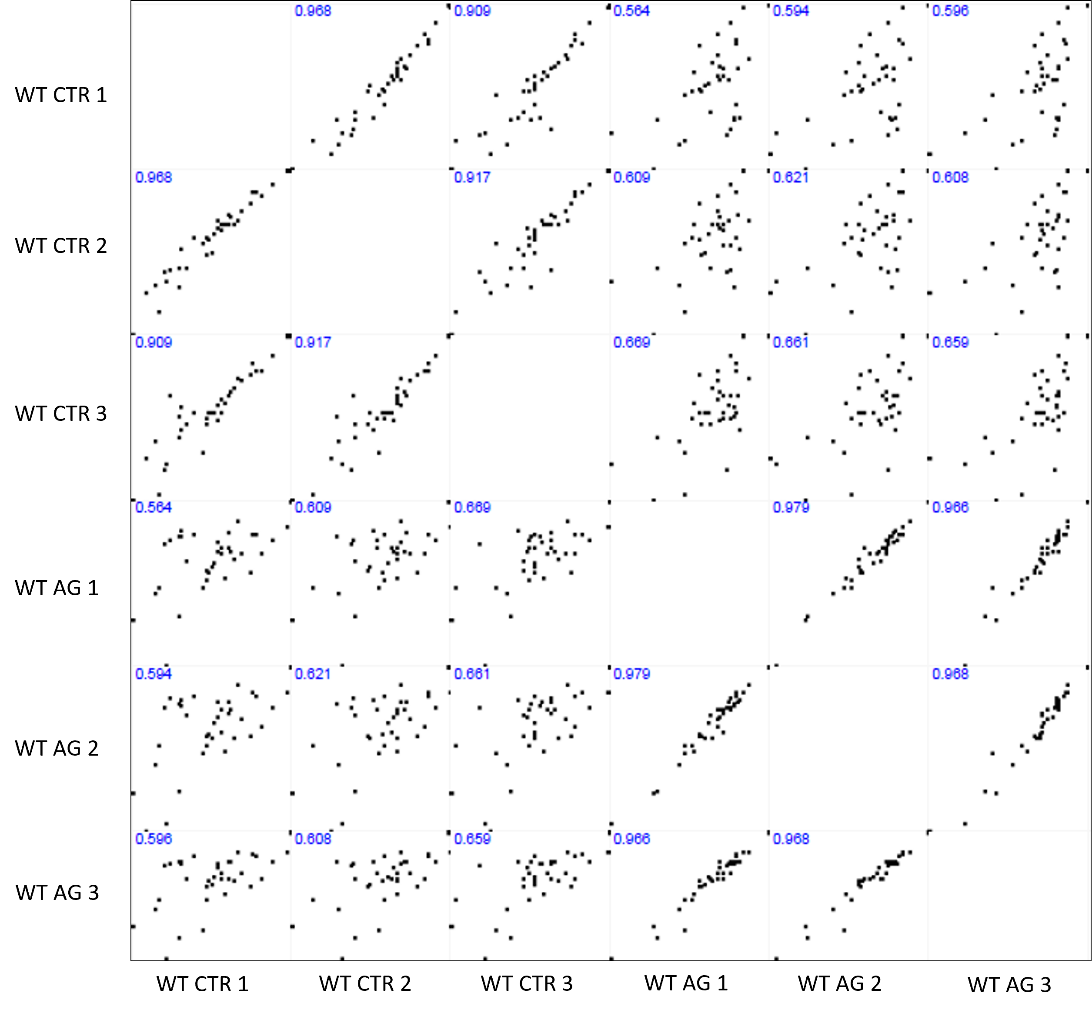

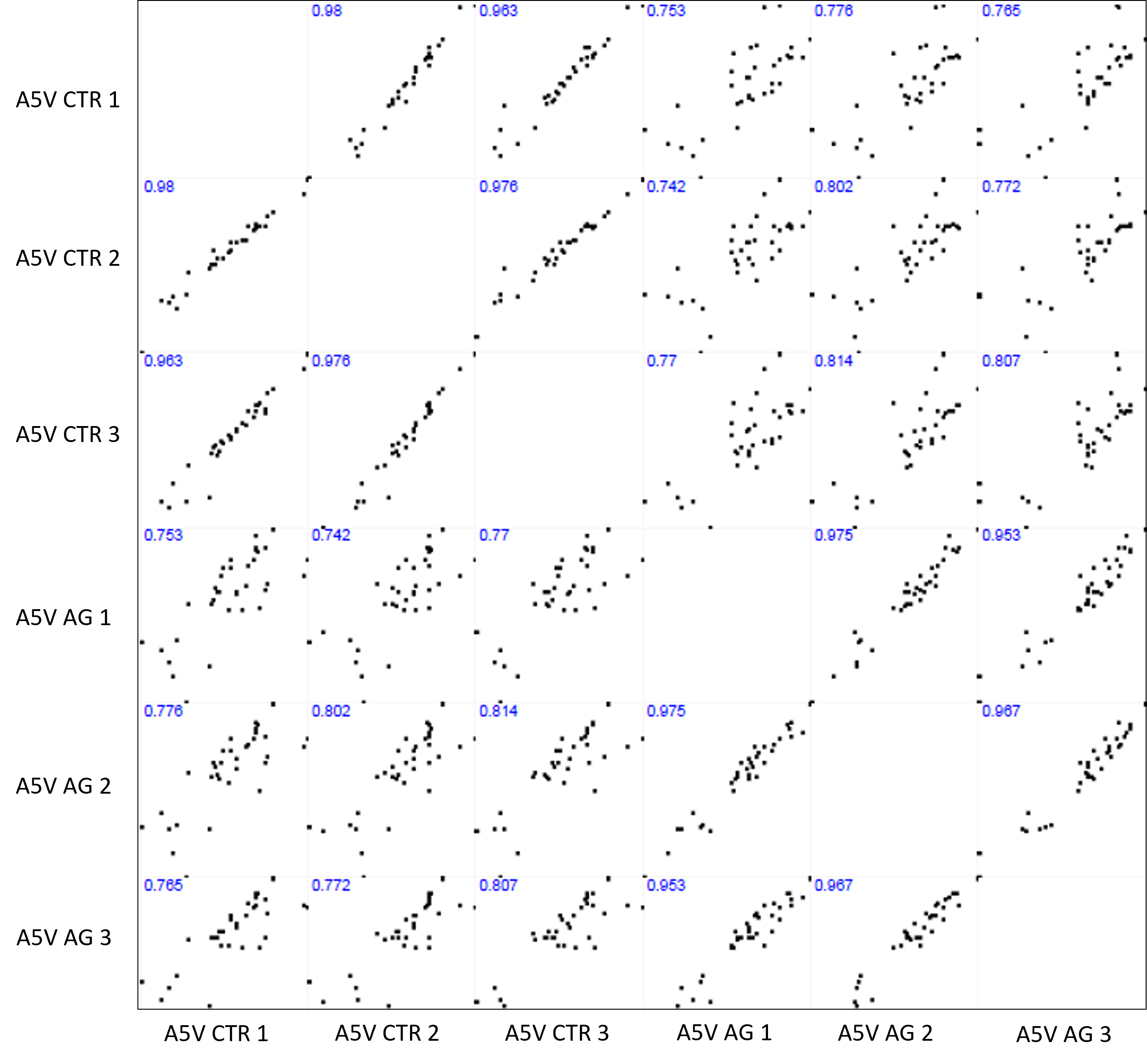

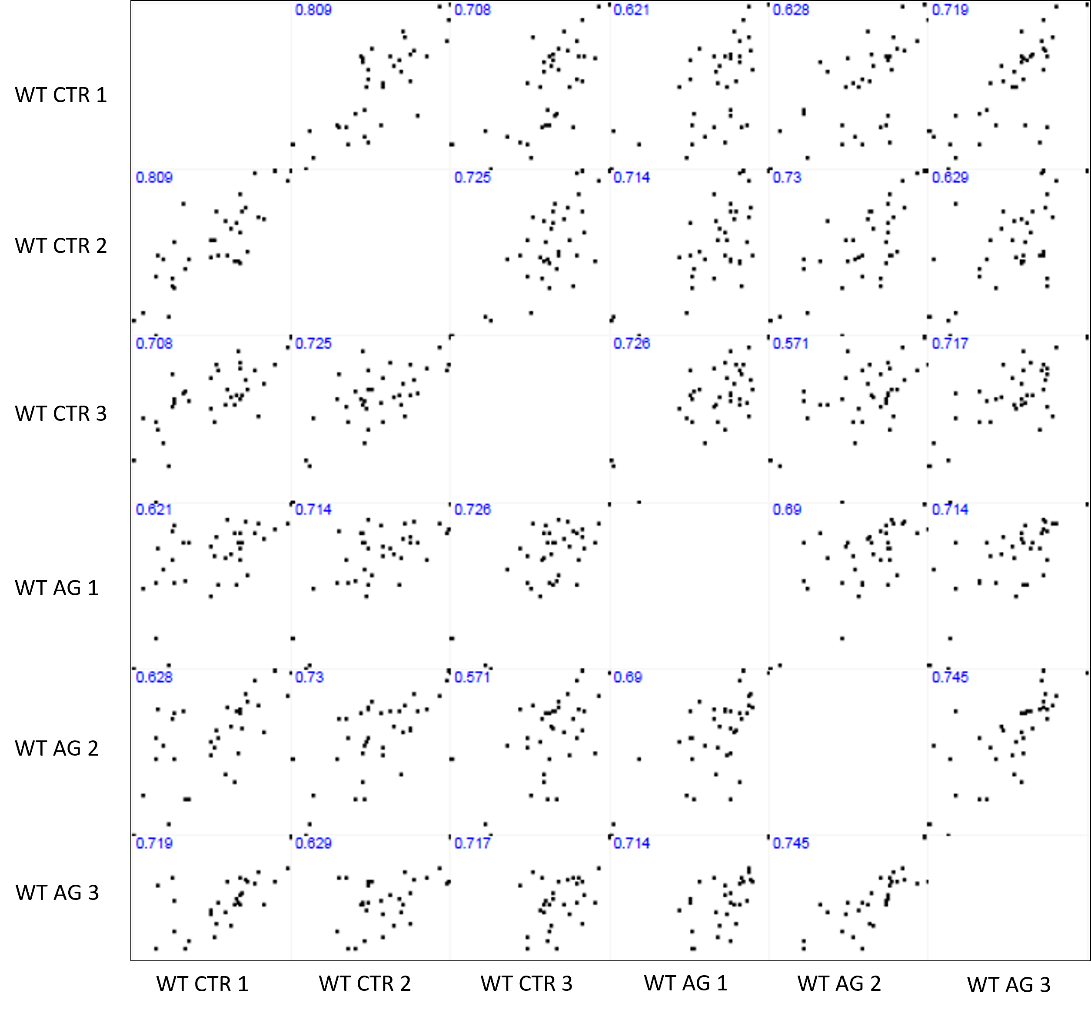

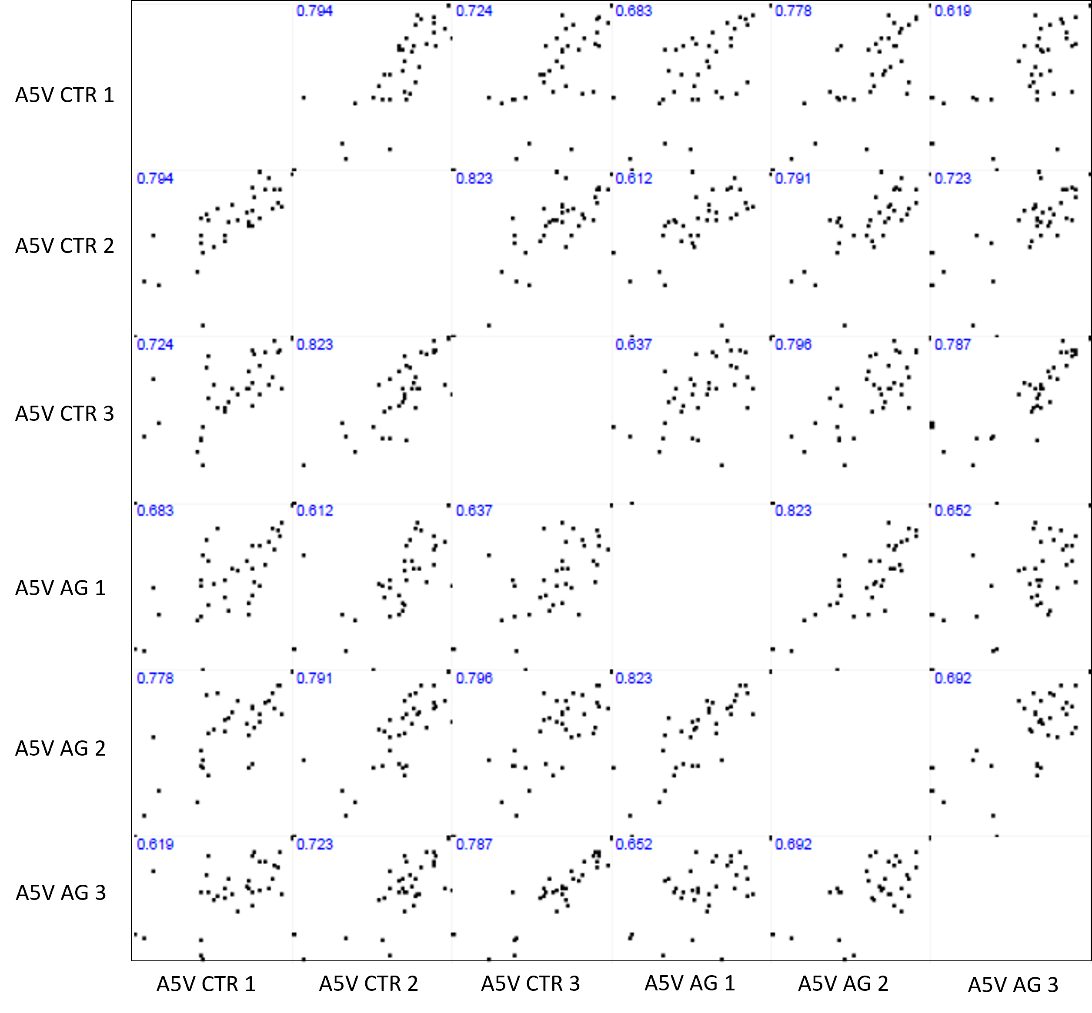

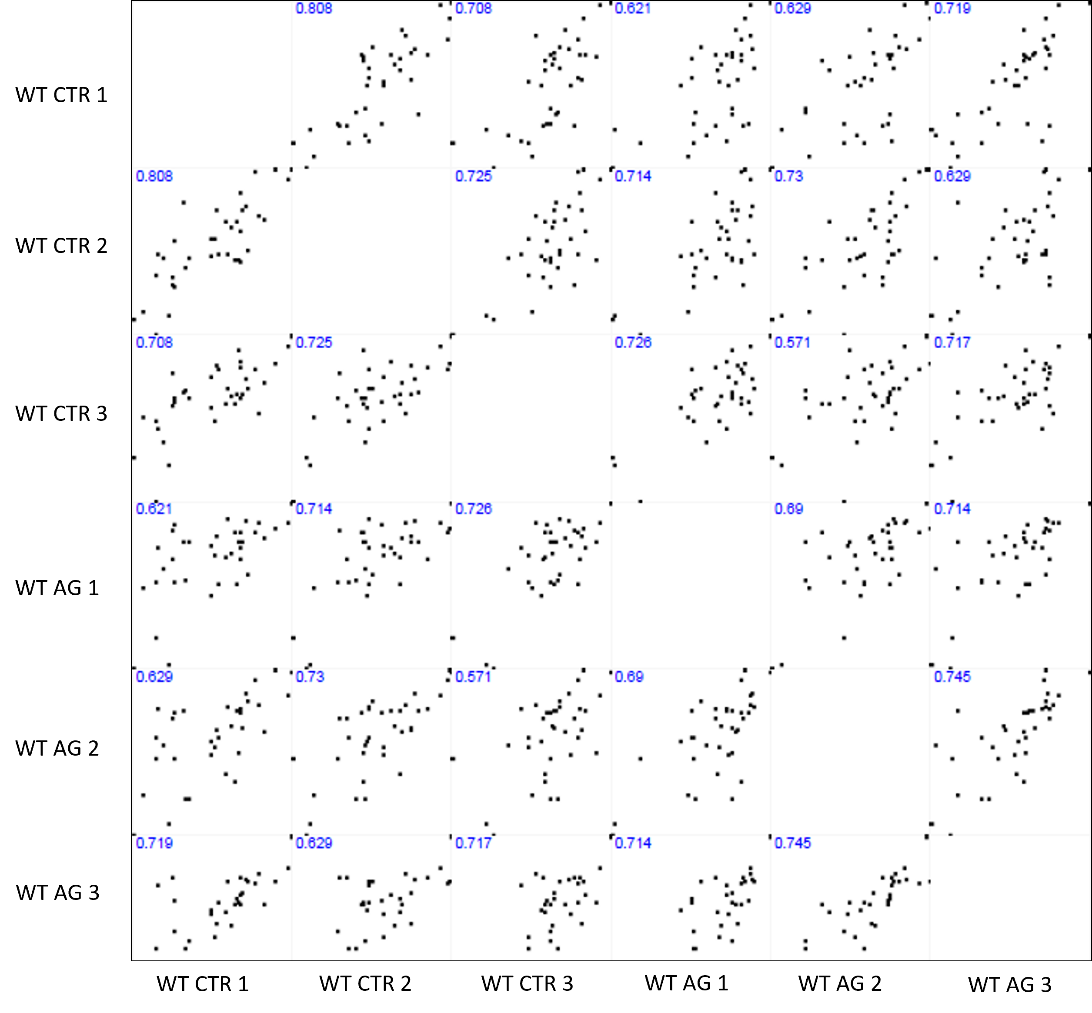

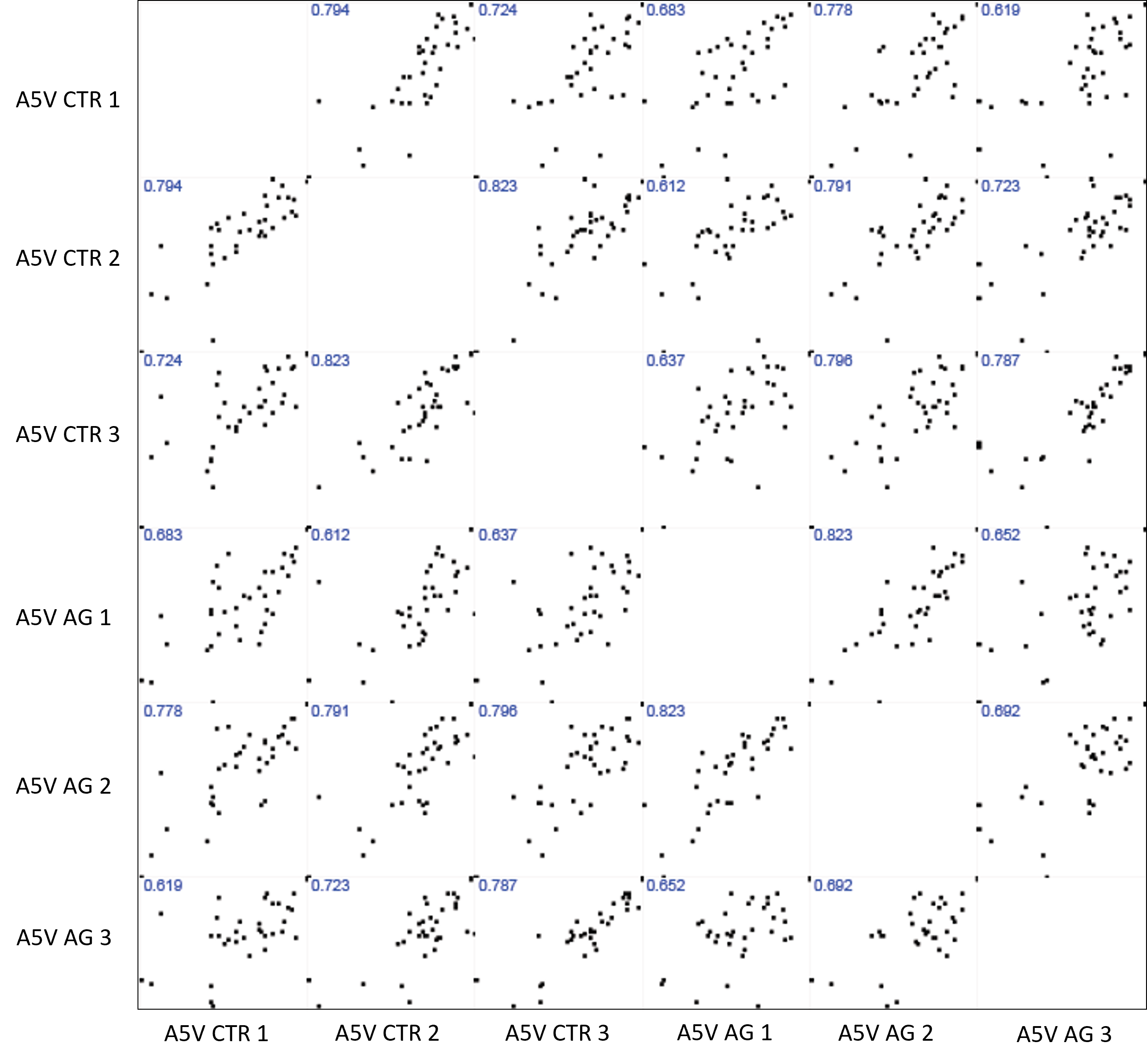

**A**

**Figure S3**. Pearson correlation between biological triplicates of SOD1 WT and A5V. (A) Raw data without using normalization by the SOD1 proteotypic peptide; (B) Raw data with normalization using SOD1 proteotypic peptide[R].TLVVHEKADDLGK.[G]; (C) Raw data with normalization using SOD1 proteotypic peptide[K].HGGPKDEER.[H]; (D) Raw data with normalization using SOD1 proteotypic peptide[K].GDGPVQGIINFEQK.[E]; (E) Raw data with normalization using SOD1 proteotypic peptide[K].ADDLGKGGNEESTK.[T].

**E**

**D**

**C**

**B**

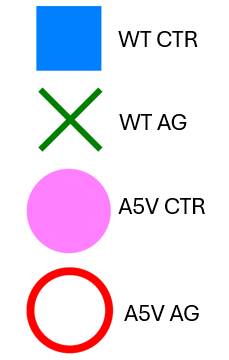

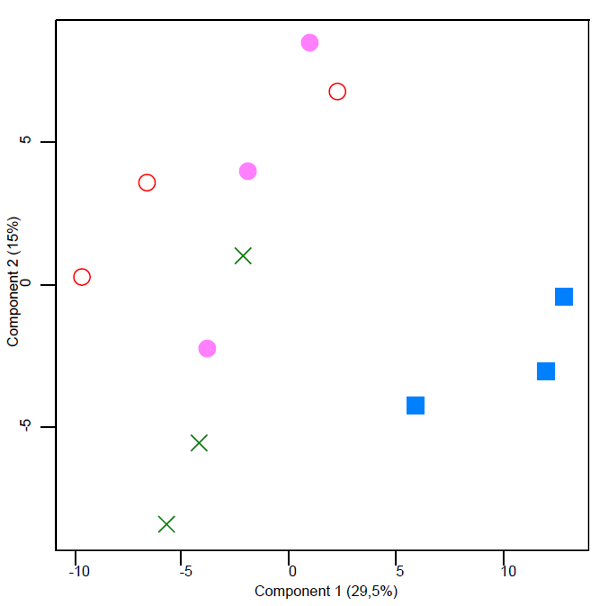

[R].TLVVHEKADDLGK.[G]

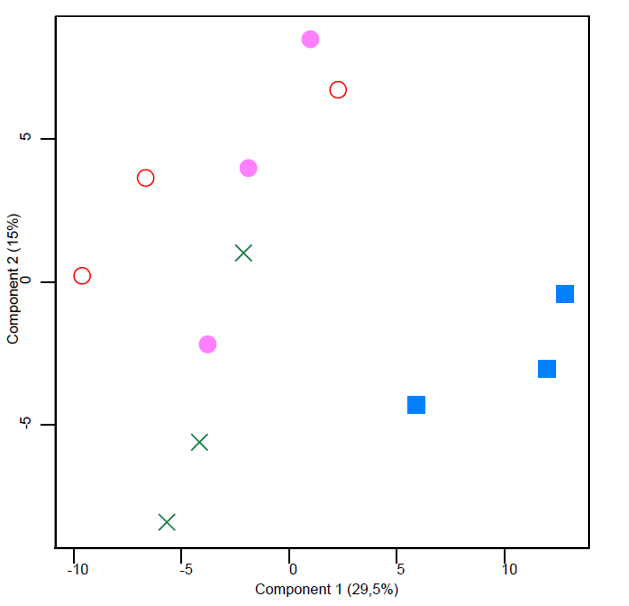

Raw Data

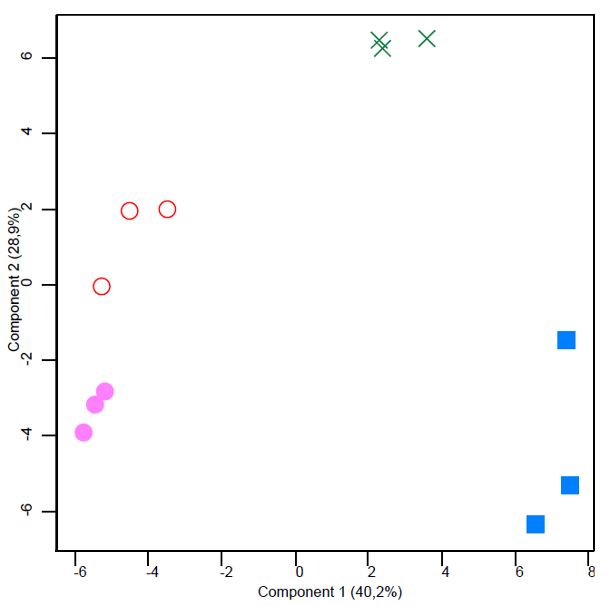

[K].HGGPKDEER.[H]

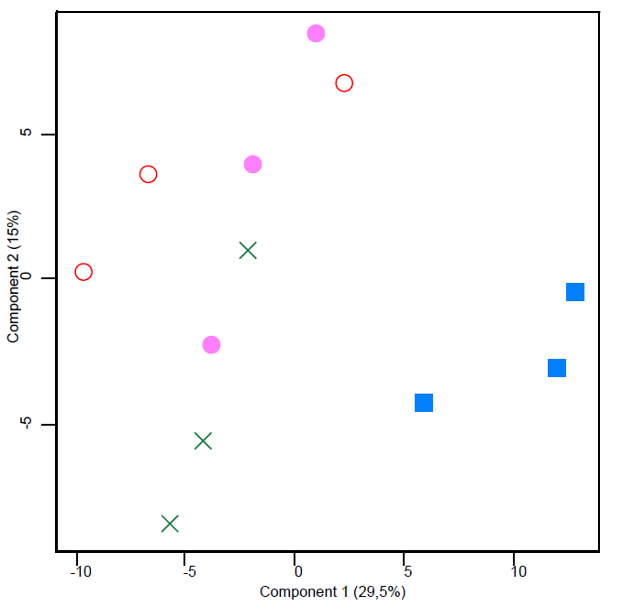

[K].GDGPVQGIINFEQK.[E]

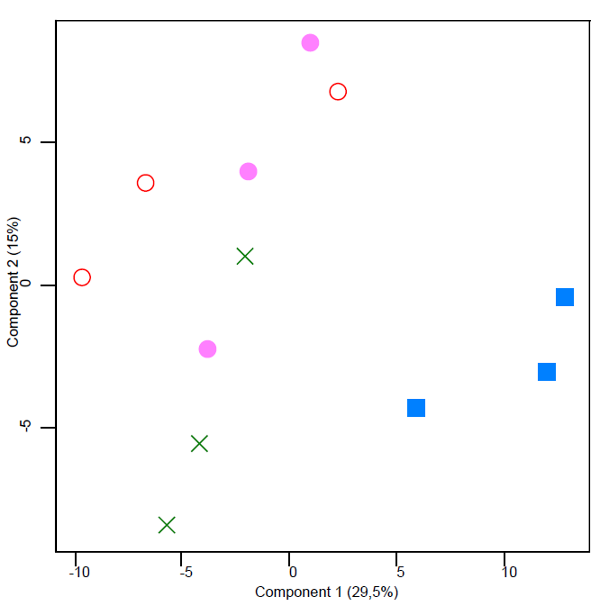

[K].ADDLGKGGNEESTK.[T]

**A**

**B**

**D**

**C**

**E**

**Figure S4**. Principal component analysis (PCA) performed to compare clustering between biological triplicates of SOD1WT and A5V, control and aging. (A) Analysis performed for raw data, without normalization; (B) Analysis performed for data normalized from the proteotypic peptide [R].TLVVHEKADDLGK.[G]; (C) Analysis performed for data normalized from the proteotypic peptide [K].HGGPKDEER.[H]; (D) Analysis performed for data normalized from the proteotypic peptide [K].GDGPVQGIINFEQK.[E]; (E) Analysis performed for data normalized from the proteotypic peptide [K].ADDLGKGGNEESTK.[T].

Sequence

[1].MATKAVCVLKGDGPVQGIINFEQK.[24]

[5].AVCVLKGDGPVQGIINFEQK.[24]

[11].GDGPVQGIINFEQK.[24]

[11].GDGPVQGIINFEQKESNGPVK.[31]

[25].ESNGPVKVWGSIK.[37]

[32].VWGSIKGLTEGLHGFHVHEFGDNTAGCTSAGPHFNPLSR.[70]

[38].GLTEGLHGFHVHEFGDNTAGCTSAGPHFNPLSR.[70]

[71].KHGGPKDEER.[80]

[72].HGGPKDEER.[80]

[77].DEERHVGDLGNVTADKDGVADVSIEDSVISLSGDHCIIGR.[116]

[81].HVGDLGNVTADK.[92]

[81].HVGDLGNVTADKDGVADVSIEDSVISLSGDHCIIGR.[116]

[81].HVGDLGNVTADKDGVADVSIEDSVISLSGDHCIIGRTLVVHEK.[123]

[93].DGVADVSIEDSVISLSGDHCIIGR.[116]

[117].TLVVHEKADDLGK.[129]

[117].TLVVHEKADDLGKGGNEESTK.[137]

[124].ADDLGKGGNEESTK.[137]

[124].ADDLGKGGNEESTKTGNAGSR.[144]

[130].GGNEESTKTGNAGSR.[144]

**Table S1**. SOD1 peptide sequence formed during trypsinization.
